# Computational design of soluble functional analogues of integral membrane proteins

**DOI:** 10.1101/2023.05.09.540044

**Authors:** Casper A. Goverde, Martin Pacesa, Nicolas Goldbach, Lars J. Dornfeld, Petra E. M. Balbi, Sandrine Georgeon, Stéphane Rosset, Srajan Kapoor, Jagrity Choudhury, Justas Dauparas, Christian Schellhaas, Simon Kozlov, David Baker, Sergey Ovchinnikov, Alex J. Vecchio, Bruno E. Correia

**Affiliations:** Laboratory of Protein Design and Immunoengineering, École Polytechnique Fédérale de Lausanne and Swiss Institute of Bioinformatics; Lausanne, Switzerland; Department of Structural Biology, University at Buffalo, Buffalo, NY, USA; Department of Biochemistry, University of Washington; Seattle, WA, USA; Institute for Protein Design, University of Washington; Seattle, WA, USA; John Harvard Distinguished Science Fellowship Program, Harvard University; Cambridge, MA, 02138 USA; Howard Hughes Medical Institute, University of Washington; Seattle, WA, USA

## Abstract

*De novo* design of complex protein folds using solely computational means remains a significant challenge. Here, we use a robust deep learning pipeline to design complex folds and soluble analogues of integral membrane proteins. Unique membrane topologies, such as those from GPCRs, are not found in the soluble proteome and we demonstrate that their structural features can be recapitulated in solution. Biophysical analyses reveal high thermal stability of the designs and experimental structures show remarkable design accuracy. The soluble analogues were functionalized with native structural motifs, standing as a proof-of-concept for bringing membrane protein functions to the soluble proteome, potentially enabling new approaches in drug discovery. In summary, we designed complex protein topologies and enriched them with functionalities from membrane proteins, with high experimental success rates, leading to a *de facto* expansion of the functional soluble fold space.

## Introduction

Protein design enables the expansion of nature’s molecular machinery, creating synthetic proteins with novel functionalities. Traditionally, protein design has been dominated by physics-based approaches, such as Rosetta^1^. However, these methods require parametric and symmetric restraints to guide the design process, and often extensive experimental screening and optimization. This proves problematic for the design of functional proteins with complex structural topologies. Recently, structure prediction pipelines, such as AlphaFold2 (AF2)^2^ and RoseTTAfold (RF)^3^ achieved unprecedented accuracy in predicting protein structure given the amino acid sequence. With the rise of deep learning-based methods, exploring novel sequence space has become increasingly feasible, allowing the discovery of proteins with stable topologies and new functions. Deep learning powered methods have been impactful in a number of tasks that include the generation of novel designable backbones^4–8^, oligomeric protein assemblies^9,10^, proteins with embedded functional motifs^11^, novel protein structural descriptors^12^, the sequence design problem^13–15^, and more recently the generation of a diverse range of protein topologies using diffusion models^5,10,16–19^. It was also found that structure prediction networks can be inverted and used for protein design, resulting in the generation of plausible protein backbones^7,8,20^.

Nevertheless, designing protein folds with complex structures, including non-local topologies and large sizes, remains challenging yet essential for creating new protein functions. In addition to design proficiency, many questions about the fundamental determinants of protein structure and folding remain elusive, particularly regarding the generalizability of deep learning methods beyond natural protein structures and sequences. To probe some of these questions, we analyzed the protein fold space in the SCOP database^21^ and observed a segregation at the structural level between proteins in the soluble proteome and in the cell membrane environments (Fig. 1a). We observed that 1075 membrane proteins exhibit unique topologies that are not found in soluble form, with only 189 folds being present in both soluble and membrane environments. This raises the question whether integral membrane protein topologies have some fundamental structural features that preclude them from existing in the soluble fold space. Consequently, we investigated if membrane folds could be designed as soluble analogs, thus achieving a *de facto* fold expansion of the soluble proteome and creating opportunities for designing novel functions using these previously inaccessible protein folds. While there has been previous work on the solubilization of near-native membrane proteins using physics-based and empirical methods^22–26^, no generalisable approach for the computational design of soluble membrane topologies with preserved functional aspects has been devised.

**Figure 1.**
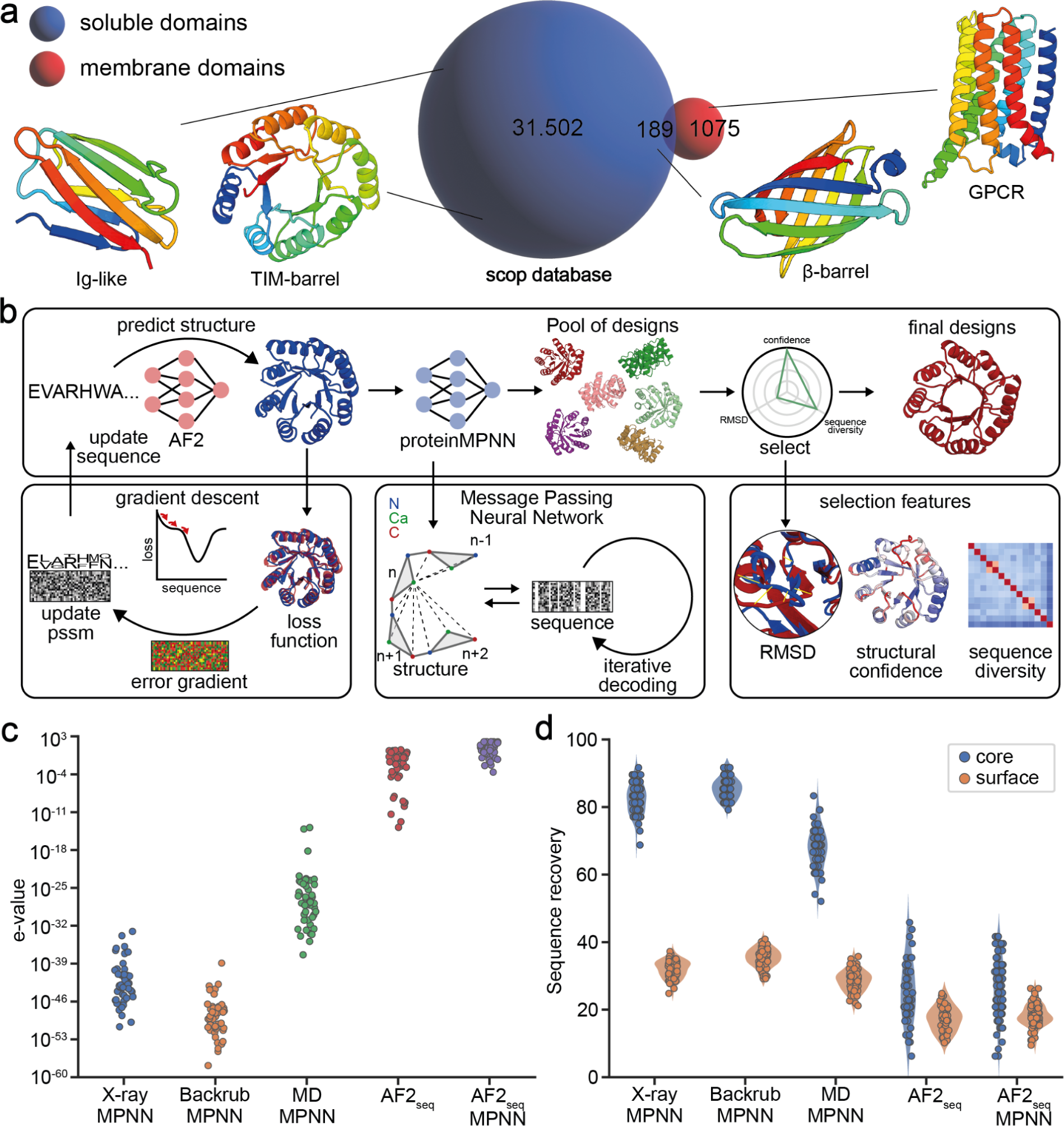
| Overview of the fold space across different environments and computational design approach. **a,** Overview of the occurrence of soluble and membrane folds in the SCOP structural database with depictions of selected representatives. **b,** Schematic representation of the integrated design pipeline for backbone and sequence generation. Given a target structure, an initial sequence is generated using AF2 through loss function optimization. The resulting structure is then passed to ProteinMPNN to sample novel sequences for a given fold. ProteinMPNN designs are filtered based on structural similarity to the target, confidence and sequence diversity., Novelty of generated sequences resulting from different backbone sampling methods evaluated by e-values relative to the non-redundant protein sequence database. **d,** Sequence recovery of core and surface residues of TIM-barrel ProteinMPNN designs generated based on the reference crystal X-ray structure (5BVL), Rosetta-perturbed backbones (backrub protocol), MD simulation trajectories, or AF2_seq_ generated structures.

To this end, we developed a computational pipeline for robust *de novo* protein design based on the inversion of the AF2 network^8^ coupled with sequence design using ProteinMPNN^15^ (Fig. 1b). Our approach allowed us to computationally design highly stable folds that were previously very challenging (Ig-like folds, β-barrels, TIM-barrels), as well as soluble analogues of integral membrane protein folds (claudin, rhomboid protease, G-protein coupled receptor (GPCR)) without the need for parametric design restraints or subsequent experimental optimization. Finally, we demonstrate that the soluble analogues can be designed in a conformation-specific manner while preserving native functional motifs with structurally elaborate features and of biological and therapeutic relevance, such as G-protein binding interfaces and toxin-receptor interaction sites. Our findings showcase the remarkable success and accuracy of deep learning-based methods in protein design, paving the way for exploring new protein topologies and sequences for improved functional design strategies.

## Results

### Structure-driven sequence generation using deep learning

AF2-based design approaches have been shown to generate plausible protein backbones^8,27,28^, however, their performance in sequence design was suboptimal as evidenced by the low experimental success rates^8,9,15^. Wicky and coworkers^9^ have demonstrated the efficiency of using ProteinMPNN on AF2-generated structures to enhance their expression and solubility, but it remained unclear whether it could be successfully applied to explore the sequence space of complex protein folds with intricate topological features including those only found in membrane environments (Fig 1a). To address this challenge, we integrated a previously developed AF2-based design approach (AF2_seq_)^8^ with the ProteinMPNN framework (Fig 1b).

In this pipeline we used AF2_seq_ to generate sequences that adopt a desired target fold. AF2_seq_ optimizes a sequence based on a loss function that comprises both topological and structural confidence loss components (see Methods), until a sequence is found that folds to the desired topology. We then apply ProteinMPNN sequence optimization to the AF2_seq_-generated starting topologies. Finally, the structures of all resulting sequences are repredicted with AF2, and filtered based on their structural similarity to the target topology (TM-score > 0.8), confidence scores (pLDDT > 80), and sequence novelty relative to naturally occurring sequences (e-value > 0.1).

*In silico* assessment showed that, despite the restricted structural diversity (Extended Data Fig. 1), AF2_seq_-designed backbones enable ProteinMPNN to generate a much larger protein sequence diversity for a desired fold than classical backbone sampling methods, such as Rosetta Backrub^29^, molecular dynamics (MD) simulations (Fig. 1c and Extended Data Fig. 1). To investigate the source of the diversity, we examined sequence conservation at the core and surface of the designs following ProteinMPNN optimization, as it was originally reported to consistently recover ∼50% of the starting sequence^15^. Sequence optimisation using ProteinMPNN alone resulted in high sequence recoveries in the core of the designs, relative to the starting sequence (Fig 1d). AF2_seq_-generated designs exhibited low sequence recoveries in both the core and the surface when compared to the sequence of the target protein. This indicates that the novelty and designability of our backbones primarily stem from the novel backbone templates generated by AF2_seq_. Interestingly, increasing levels of Gaussian noise applied to the backbone prior to ProteinMPNN sequence design could also reduce sequence recovery (Extended Data Fig. 2a), however, at the expense of low-confidence predictions which deviate significantly from the target fold (Extended Data Fig. 2b-d). Additionally, we found that for some more stringent design examples, the target structure could not be predicted in single sequence mode by AF2 upon ProteinMPNN redesign. However, when using a combination of AF2_seq_ and ProteinMPNN (AF2_seq_-MPNN), we found the input sequence to result in accurate structural predictions of the target folds (Extended Data Fig. 3). Therefore, we sought to test whether our design strategy would be successful in designing protein folds that have thus far been challenging to other approaches.

### Computational design of topologically complex protein folds

To identify challenging design targets for our pipeline, we quantified the topological complexity of protein folds using metrics of protein length and sequence contact order (Extended Data Fig. 4, Methods). Based on this assessment and on how challenging some folds have been for computational design, we selected three folds to test our approach: Ig-like fold (IGF), β-barrel fold (BBF), and TIM-barrel fold (TBF) (Fig. 2a). The IGF is one of the most prevalent folds in nature and is an essential building block of immunological effectors and therapeutics, such as antibodies and receptors^30–32^. The IGF consists of two stacked β-sheets, presenting a significant design challenge. This is due to its non-local interactions and susceptibility to aggregation via edge β-strands^33,34^, previously requiring strict parametric and symmetry restraints during design^35,36^. Using our AF2_seq_-MPNN protocol (Fig. 1b), we designed novel IGFs that are significantly distant from natural protein sequences (Fig. 2b). We selected 19 designs for experimental characterisation based on AF2 confidence scores and sequence diversity (Extended Data Fig. 5-6). Seven designs were soluble, with 4 designs exhibiting monodisperse peaks in solution (Extended Data Fig. 7). Exemplified by IGF_10 (Fig. 2c), the designed IGFs exhibited the typical β-sheet rich secondary structure profile according to circular dichroism (CD) spectroscopy, along with unusually high thermostability (Fig. 2c, Extended Data Fig. 7)^37^.

**Figure 2.**
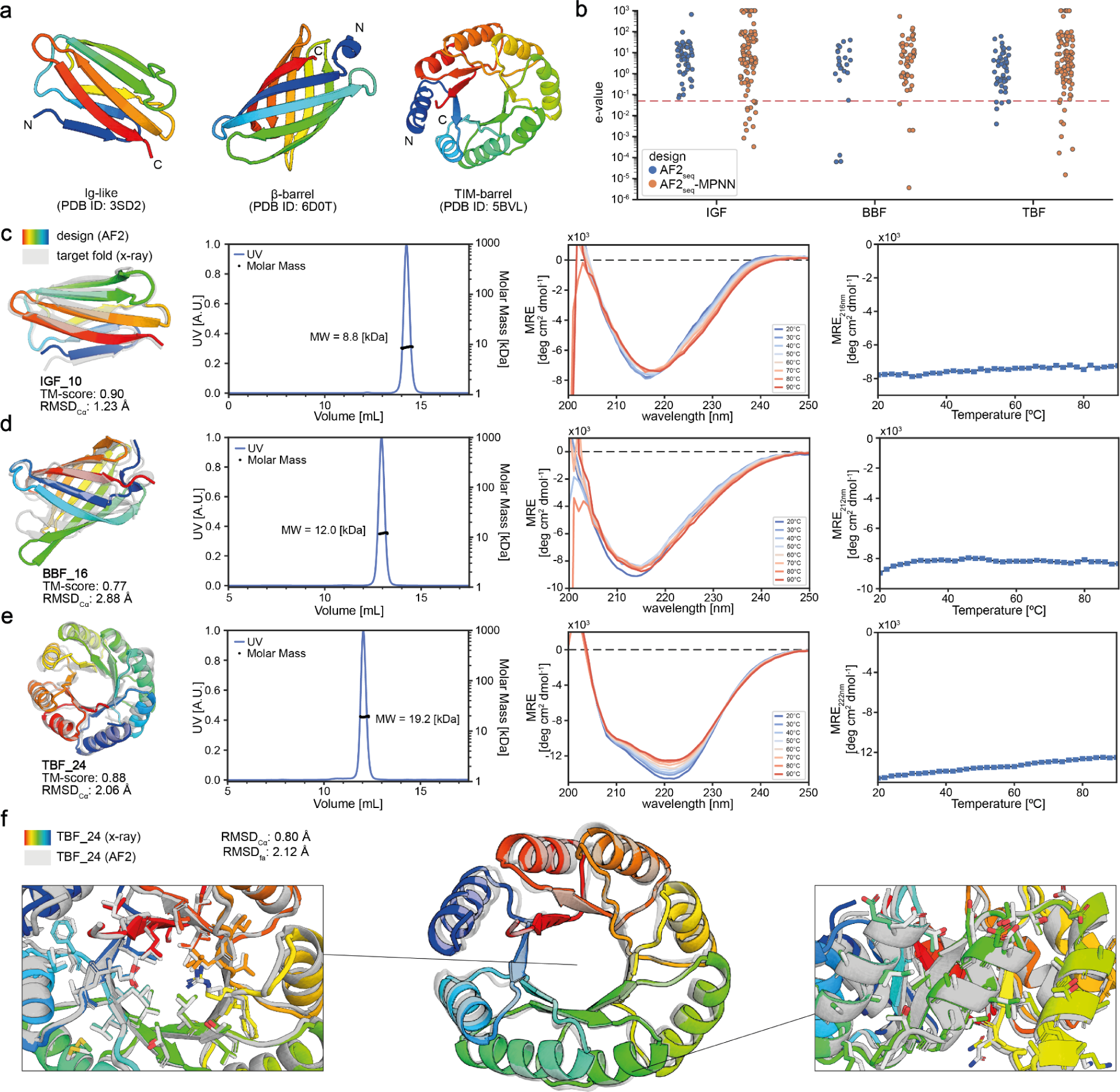
| Experimental characterization of designed complex protein topologies. **a,** Cartoon depiction of three protein topologies which have been challenging for computational design: Ig-like Fold (IGF), β-barrel Fold (BBF), and TIM-barrel Fold (TBF). **b,** Closest e-value hit of the AF2_seq_ and AF2_seq_-MPNN designs when searching a non-redundant protein sequence database. Significance threshold of 0.05 is highlighted, indicating little sequence homology to natural sequences. Characterization of design **c,** IGF_10, **d,** BBF_16, and **e,** TBF_24, respectively. Superposition of the design (color) and the target fold (gray), the corresponding SEC-MALS measurement, CD-spectra at different incubation temperatures, and the CD melting curve. **f,** X-ray structure of the TIM-barrel fold TBF_24 (colored) superimposed on the design model (gray). RMSD_Cα_ - root mean square deviation computed over the Cα atoms of the backbone. RMSD_fa_ - root mean square deviation computed over all the atoms in the structure.

Next, we attempted to design a *de novo* β-barrel (BBF): a fold present both in the soluble and membrane proteome, with applications as small molecule binders, transporters, and sensors^38–42^. It consists of eight antiparallel β-strands with precise hydrogen bonding patterns^38^, making its design extremely challenging. Previously, Dou et al. utilized a set of design principles involving the introduction of glycine kinks, β-bulges, and tryptophan corners to alleviate backbone strain and allow for continuous hydrogen bonding connectivity^38^. We investigated whether our approach could successfully design BBFs without explicitly defining such constraints. We experimentally characterized 25 designs, out of which 6 were found to be folded and monomeric in solution, while exhibiting high thermal stability (Fig 2d, Extended Data Fig. 8). Sequence analysis of the designs revealed a high glycine residue recovery at glycine kink positions (Extended Data Fig. 9), as observed by Verkuil et al^43,43^(Extended Data Fig. 9). This demonstrates that not all empirically-derived features are necessary for successful BBF design and a larger uncharted sequence space can be explored.

Finally, we attempted to design the TIM-barrel fold (TBF); a challenging protein topology of paramount importance in biology, as its structure is highly proficient in supporting enzymatic active sites, making it an ideal candidate for the design of enzymes with novel catalytic functions^44–46^. The TIM-barrel fold comprises eight parallel-paired β-strands, each separated by an alpha helix, resulting in long-range interactions between the β-strands^47^. The TBF has been a long-standing challenge in protein design^47–49^, and only very recently, several studies successfully designed this fold^10,50,51^. Previous TBF design strategies imposed symmetry and parametric restraints both at the structural and sequence level^50,51^. With our pipeline, we were able to design TBFs without any constraints, allowing for a larger structural and sequence diversity, and even asymmetry, which could potentially accommodate more complex enzymatic sites (Extended Data Fig. 10). We experimentally assessed 25 designs, 5 of which were monomeric, folded and highly thermostable in solution (Fig. 2e, Extended Data Fig. 11). To confirm the accuracy of our design, we solved a crystal structure of TBF_24 at 1.34 Å resolution (Fig. 2f). Our asymmetric design showed noticeable structural deviations from the initial symmetric template (Fig. 2f), with an overall backbone RMSD_Cα_ of 2.06 Å (Extended Data Fig. 12). When comparing the x-ray structure to the designed model, the Cα and full-atom RMSD_fa_ are 0.80 Å and 2.12 Å, respectively (Fig. 2f). These structural comparisons demonstrate the remarkable accuracy of our design approach, further underlined by the almost identical sidechain placement within both the core and peripheral regions of the protein (Fig. 2f). Given the encouraging results from our initial designs, we wondered whether our approach would allow us to probe the sequence space of topologies not present in the soluble proteome, such as those of integral membrane proteins.

### Expanding the soluble fold space with membrane protein topologies

In a domain analysis performed over the SCOP database, we observed that both the soluble and membrane proteome each encompass a group of unique structural protein topologies, with only a narrow overlap between the two (Fig. 1a, Fig. 3a). This prompted us to ask whether it is possible to design soluble analogues of such membrane-only folds or whether they contain intrinsic structural features that preclude them from existing in soluble form. Previous studies have demonstrated that simply substituting exposed hydrophobic residues for polar or charged amino acids might not be sufficient to solubilize them, since the interactions between the surface residues have to be carefully considered^22,52–54^. To address this question, we set out to design soluble analogues of membrane proteins using the AF2_seq_-MPNN pipeline (see Methods). We selected three membrane folds to test the design strategy: the claudin fold^55^, the rhomboid protease fold^56^, and the GPCR fold^57^ (Fig. 3b).

**Figure 3.**
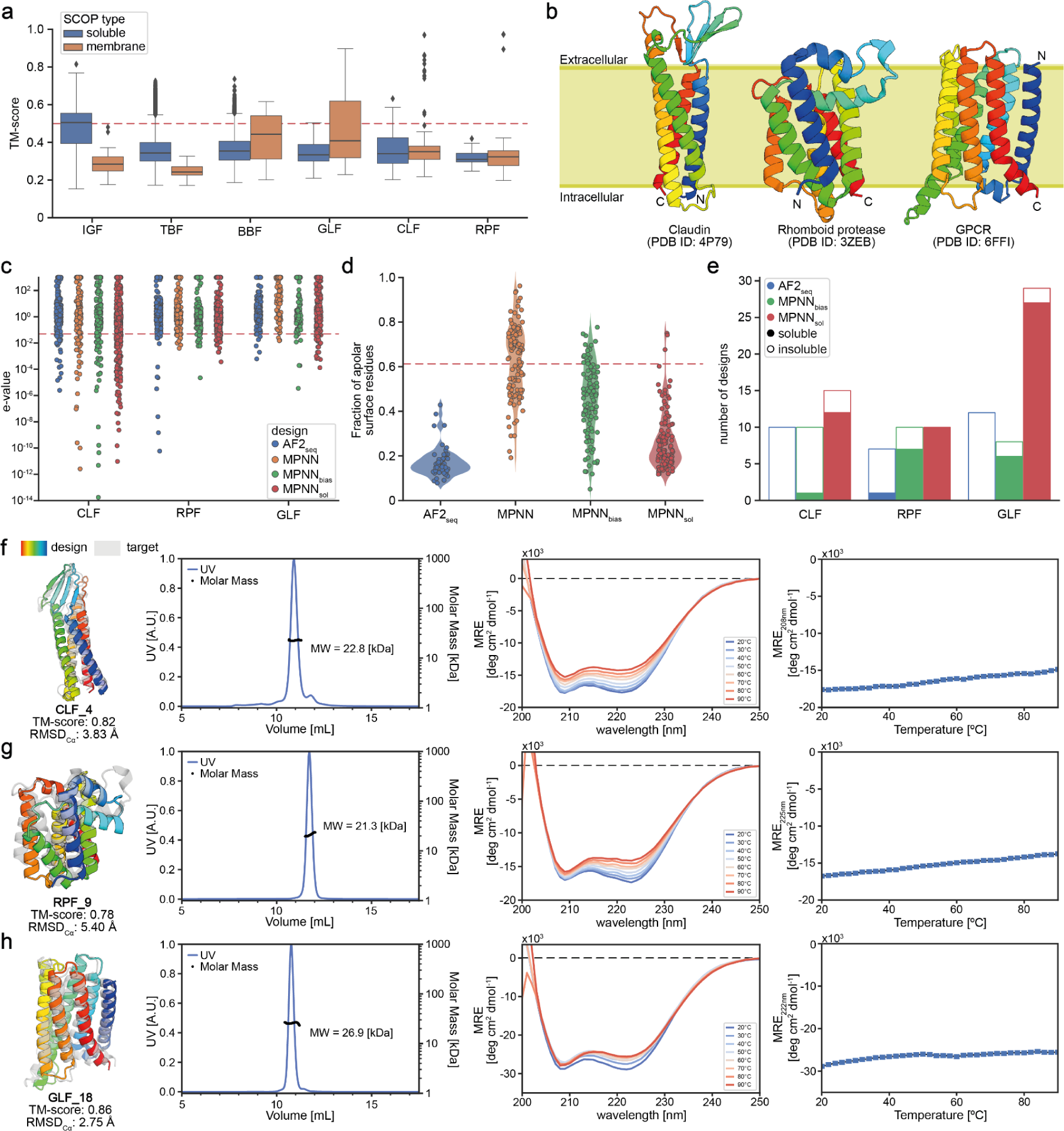
| Experimental characterization of soluble analogues of membrane proteins. **a,** Structural similarity for each of the target folds against the SCOP database. TM-score cut-off of 0.5 is highlighted, denoting significant structural similarity as compared to the reference fold. **b,** Cartoon representation of three transmembrane topologies chosen to be redesigned as soluble folds: Claudin-Like Fold (CLF), Rhomboid Protease Fold (RPF) and GPCR-Like Fold (GLF). **c,** Closest e-value hits of the solubilized CLF, RPF and GLF against a non-redundant protein sequence database. The vast majority of the designed sequences differ substantially from natural sequences as indicated by e-values higher than the significance threshold of 0.05 (red line). **d,** Fraction of hydrophobic residues found on the surface of the GLF designs using different sequence-generation methods following AF2_seq_ backbone generation. The fraction of surface hydrophobics of the native GLF is 0.61 (red line) **e,** Number of designs resulting in soluble expression of the designed soluble membrane protein analogues. Experimental characterization of **f,** CLF_4, **g,** RPF_9 and **h,** GLF_18. Comparison between the design (color) and target fold (gray), solution behavior by SEC-MALS, CD spectra at different incubation temperatures and melting temperature profiles by CD.

Initial designs by AF2_seq_ exhibited a high sequence novelty when compared to natural proteins (Fig. 3c) and a low fraction of surface hydrophobics (Fig. 3d), however, none could be expressed in soluble form (Fig. 3e). We attempted to optimize the sequences using the standard ProteinMPNN model, but the resulting sequences consistently recovered the surface hydrophobics, most likely due to the topology’s similarity to membrane proteins encountered during training (Fig. 3d). Biasing the amino acid sampling towards hydrophilic amino acids (AF2_seq_-MPNN_bias_) only marginally improved the solubility of the designs (Fig. 3d). Therefore, we retrained the ProteinMPNN network using a dataset of only soluble proteins, which we named soluble MPNN (MPNN_sol_) (Fig. 3c-3e). AF2_seq_-MPNN_sol_ was able to produce novel sequences with high confidence scores predicted by AF2 and low fraction of surface hydrophobics (Fig. 3d, Extended Data Fig. 5f). As a result, we were able to generate high confidence designs of membrane protein topologies that do not exist in the soluble proteome.

We started by designing soluble analogues of the claudin fold, a class of proteins involved in the formation of tight junctions, which are critical in controlling the flow of molecules between layers of epithelial and endothelial cells^58^. Claudin folds are composed of an alpha/beta mixed secondary structure where there are four transmembrane alpha helices and an extracellular β-sheet^58^. The composition of the β-sheet determines the type of tight junction between cells that is being formed, resulting in highly selective permeability of ions and solutes^58^. Claudin-targeting therapies hold great promise as novel cancer therapies and soluble claudin analogues could introduce a novel route to screen for claudin binders^59^. We tested 13 designs for the claudin-like fold (CLF) out of which 10 were found to be expressed in soluble form (Fig. 3e). Five designs were further biochemically characterized, and three were monomers in solution according to SEC-MALS and were folded, with two showing a T_m_ above 90 °C (Fig. 3f, Extended Data Fig. 13). The CLF designs showed a sequence identity below 13% relative to the native fold and nearest e-values to natural sequences below 0.063 (Fig. 3c, Extended Data Fig. 5). AF2 predicted structures in comparison with the designed models exhibited RMSDs_Cα_ ranging from 2.84 - 4.03 Å (Extended Data Fig. 13). The CLF design series showed that our approach could successfully design soluble analogues of membrane proteins with rather simple topologies such as 4-helix bundles.

Next, we attempted to design a larger fold and a more intricate topology, the rhomboid protease fold (RPF). The RPF comprises six transmembrane alpha helical domains, with many intricate loops and long-range contacts^60^ (Fig. 3b). Additionally, it harbors a serine-histidine catalytic dyad buried in the cell membrane, allowing it to cleave transmembrane protein domains, which plays an important role in cell signaling, making it a therapeutically interesting target^61,62^. We selected 15 designs for protein expression, out of which 13 were found to be soluble (Fig. 3e) and five were selected for further experimental characterization. Three of the five designs showed a single monomeric species in solution and the expected helical secondary structure as assessed by CD (Fig. 3g, Extended Data Fig. 14). All of the three monomeric species of RPFs exhibited high thermal stability, with T_m_ above 90 °C (Fig. 3g, Extended Data Fig. 14). Interestingly, the AF2 structure predictions of the designs were less accurate than those of the CLF designs, with the RMSD_Cα_s between models and predictions ranging between 3.34 and 5.57 Å (Fig. 3g, Extended Data Fig. 14). Overall, the high RMSD_Cα_s between design models and AF2 predictions highlight the inherent difficulty in designing folds with such structural complexity.

Finally, we tested our design approach in one of the most prevalent membrane folds in nature: the GPCR fold. GPCRs are the largest and most diverse family of membrane receptors in eukaryotes, playing important roles in signaling pathways^57^. About 34% of all US Food and Drug Administration (FDA) approved drugs target GPCRs and they remain the most studied drug target^63,64^. The core topology of GPCRs comprises seven transmembrane helices that facilitate numerous non-local interactions, enabling them to bind a variety of ligands, including photoreceptors, odors, pheromones, hormones, and neurotransmitters^57^. *De novo* design of GPCR-like folds (GLF) provides the potential of creating new small molecule receptors and protein scaffolds with functional sites of the GPCRs. We tested 56 designs out of which 36 were expressed to be soluble, and we selected the 10 most highly expressed designs for further biochemical characterization. From the ten designs, nine were monodisperse monomers in solution, and all showed the characteristic CD signature of alpha-helical rich proteins (Fig. 3h, Extended Data Fig. 15). All ten designs also showed high thermal stabilities (T_m_ > 90 °C) (Fig. 3h, Extended Data Fig. 15).

Next, we attempted to crystallize the soluble analogues of membrane topologies and obtained high resolution X-ray structures for one claudin, one rhomboid protease, and two GPCR designed folds (Fig. 4, Extended Data Fig. 16). The CLF_4 structure exhibited exceptional design precision in both backbone and sidechains, as indicated by the backbone RMSD_Cα_ of 0.73 Å and full-atom RMSD_fa_ of 1.28 Å (Fig. 4a). The comparison between CLF_4 and the native claudin revealed accurate secondary structural element positioning, with an RMSD_Cα_ of 3.63 Å, where most of the deviation arises from the β-sheet region (Fig. 4b, Extended Data Fig. 17). Additionally, the four helices are mostly hydrophilic, as evidenced by the low lipophilicity potential on the surface (Fig. 4c). In the case of the rhomboid protease design RPF_9, we observed high accuracy between the X-ray structure and the design model, with a backbone RMSD_Cα_ of 0.97 Å and full-atom RMSD_fa_ of 1.83 Å. However, the structural similarity was significantly lower when compared to the target fold, as indicated by the backbone RMSD_Cα_ of 5.67 Å (Fig. 4d, 4e). Specifically, large structural deviations in the first extracellular loop (Extended Data Fig. 18) were observed, which could be due to its native positioning in the water/membrane interface^61^. The designed RPF_9 fold showed significantly increased hydrophobicity on the transmembrane surface when compared to the native RPF fold (Fig. 4f). Lastly, the designed GLFs preserved the canonical seven-helical bundle characteristic of native GPCRs. In terms of structural accuracy, the crystal structures were in very good agreement with the design models, 1.05 Å and 0.88 Å of RMSD_Cα_ for GLF_18 and GLF_32, respectively (Extended Data Fig. 16, Fig. 4g, 4h). This accuracy further extended to the side-chain level, where comparisons of crystal structures vs the design models for GLF_18 and GLF_32 showed 1.54 and 1.40 Å full-atom RMSD_fa_s, respectively. Comparing the structures of the soluble analogues to that of the reference native GPCR, the overall backbone RMSD_Cα_s were 3.08 and 3.51 Å for GLF_18 and GLF_32, respectively (Extended Data Fig. 16, Fig. 4g, 4h). By analyzing the lipophilicity potential at the surface^61^ of the designed proteins we can observe a clear transition from an initially hydrophobic surface to a hydrophilic one (Fig. 4i, Extended Data Fig. 16). Additionally, much of the sequence signatures of the GPCR fold were removed, such as the evolutionary conserved DRY motif in the first intracellular loop^65^, the (N/D)PxxY motif in the seventh helix^66^ and the transmembrane proline rich domains^67,68^ (Extended Data Fig. 19). This demonstrates that by design one can explore very diverse sequence spaces while removing potential evolutionary biases. Interestingly, at the structural level we observe that the irregular local structure of the terminal helix is preserved, while the intracellular segment of the GPCR fold exhibits notable structural deviation in GLF_18 (Extended Data Fig. 20). Our results demonstrate that integral membrane folds can be successfully designed in soluble form, hinting that these topologies share many of the same designability principles and constraints of folds present in the soluble proteome.

**Fig. 4.**
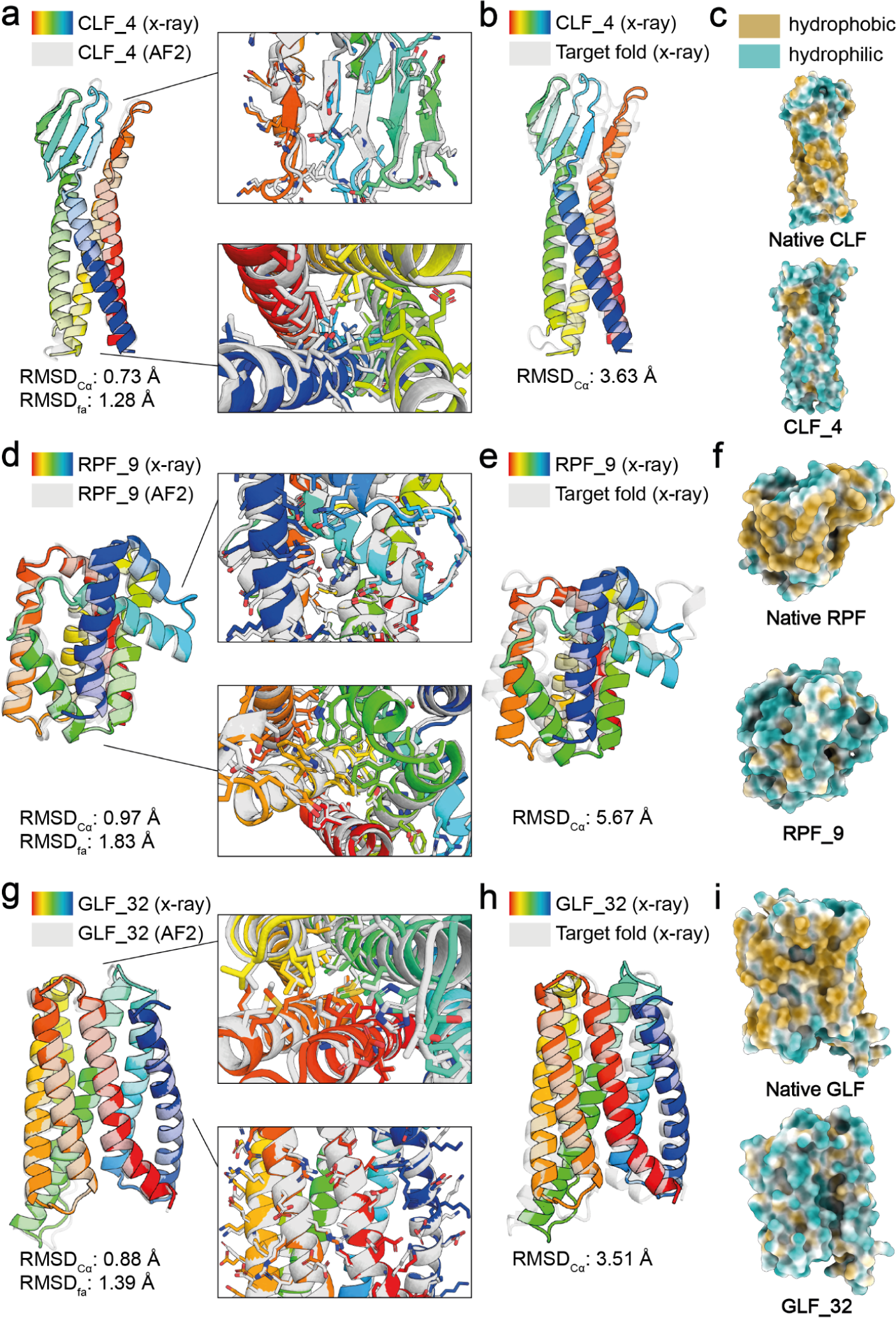
| Soluble analogues of membrane proteins solved by X-ray crystallography. **a,** X-ray structure of CLF_4 (colored) superimposed on the design model (gray). **b,** X-ray structure of CLF_4 (colored) superimposed on the design model (gray). **c,** Molecular lipophilicity potential of the surface of the claudin design target and the soluble design CLF_4. **d,** X-ray structure of RPF_9 (colored) superimposed on the design model (gray). **e,** X-ray structure of RPF_9 (colored) superimposed on the design model (gray). **f,** Molecular lipophilicity potential of the surface of the rhomboid protease design target and the soluble design RPF_9. **g,** X-ray structure of GLF_32 (colored) superimposed on the design model (gray). **h,** X-ray structure of GLF_32 (colored) superimposed on the design model (gray). **i,** Molecular lipophilicity potential of the surface of the GPCR design target and the soluble GLF_32 design. After redesign of the original membrane folds with MPNN_sol_, the hydrophobicity (yellow) of the surface is significantly reduced, and polarity increased (blue). RMSD_Cα_ - root mean square deviation computed over the Cα atoms of the backbone. RMSD_fa_ - root mean square deviation computed over all the atoms in the structure.

### Functionalization of soluble membrane protein analogues

After validating the structural accuracy of our designs, we explored the possibility of functionalizing the designed soluble analogues. To this end, we devised an approach where we explicitly fix structural segments and amino acid identities of the functional motifs during design, while the transmembrane segment is solubilised in their context (Fig. 5). We applied this strategy to the design of soluble analogues of human claudin-1 and claudin-4^69^, where varying levels of the two extracellular segments are preserved (Methods). To verify structure and function, we tested their binding to *Clostridium perfringens* enterotoxin (CpE), a common food-borne pathogen to humans known to bind claudin-1 and claudin-4 differentially^69^. Binding assays using bio-layer interferometry (BLI) indicate that soluble claudin-1 and claudin-4 exhibit binding kinetics and affinities for CpE that are comparable to their membrane-bound counterparts^69^ (Fig. 5b-5e). The claudin-1 designs exhibited lower binding affinity for CpE versus claudin-4, owing to the latter being a high-affinity CpE receptor. Notably, the higher proportion of native sequence preserved in CLN1_14 resulted in a reduced melting temperature, compared to CLN1_18 (Extended Data Fig. 21).

**Fig. 5.**
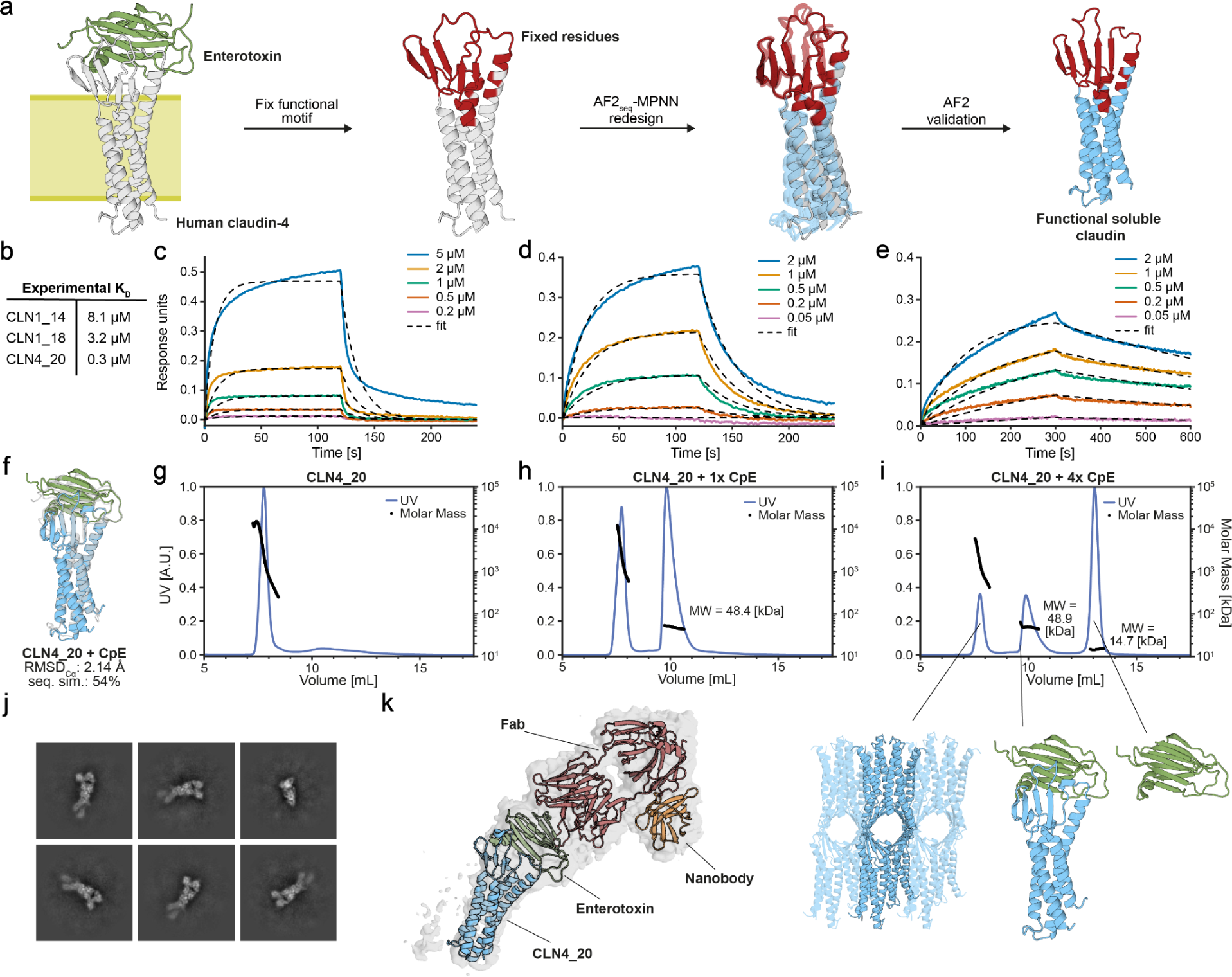
| Functionalization of soluble analogues of claudin proteins. **a,** Design workflow for solubilizing claudins with fixed functional residues. *C. perfringens* enterotoxin (CpE) is known to bind to human claudin-1 (CLN1) and claudin-4 (CLN4). **b,** Binding affinities of solubilised claudins to CpE derived from kinetic measurements. Binding kinetics of solubilized claudins **c,** CLN1_14, **d,** CLN1_18, and **e,** CLN4_20 to CpE. Association and dissociation during BLI are shown in solid lines and the respective fits as dashed lines. **f,** Cartoon depiction of design model of CLN4_20 bound to cCpE toxin (coloured) overlaid with the target fold (gray). SEC-MALS analysis of CLN4_20 mixed with **g,** 0-**h,** 1x-**i,** and 4x-molar excess of CpE toxin. **j,** Representative 2D classes of CLN4_20 bound to cCpE toxin, COP2 Fab, and a nanobody. **k,** Docked model of CLN4_20 complex into reconstructed cryoEM density. **l,** Structural overlay of the refined CLN4_20 cryoEM mode (coloured) and the design model (gray). **m,** Structural overlay of the refined CLN4_20 cryoEM mode (coloured) and the human CLN4 structure (gray).

Interestingly, we observed that CLN4_20 assembles into soluble high molecular weight oligomeric species, according to SEC-MALS (Fig. 5f-5g). The oligomeric assemblies could be disrupted by the addition of the C-terminal claudin-binding domain of CpE (cCpE) (Fig 5h-5g). This is analogous to the disassembly of high order claudin oligomers within tight junctions by cCpE in the gut^69–71^. To confirm that the soluble analogue engages the toxin in the same manner as the natural membrane-bound claudin-4, we reconstituted the complex together with a Fab and nanobody to increase its size, and determined its structure using cryoEM (Fig. 5j-5k, Extended Data Fig. 21e). We observe that both the claudin topology and the toxin binding mode are comparable to the natural complex^72^. These results suggest that the designed soluble membrane protein analogs can accommodate natural sequences and functional motifs in native-like conformations, and that certain mechanistic aspects can be recapitulated in solution.

To embed function in the soluble GPCR analogues, we applied two distinct design approaches. Firstly, we created chimeric proteins from GLFs and the intracellular loop 3 (ICL3) of the ghrelin receptor^73^ (Fig. 6a). The residues grafted from ICL3 connect TM5 and TM6 and form hydrophobic interactions with the α subunit of G_i_ in the activated ghrelin receptor^74^. The GLF-Ghrelin chimeras (GGC) were generated using a sequence transplant of the natural epitope into the corresponding region of the GLF scaffold (Fig. 6a). Using a pull-down assay, we pre-screened 16 chimeric designs for binding and found that 9 designs bind to the ICL3-targeting antibody^73^, while GLF scaffolds without the ICL3 did not show any binding (Extended Data Fig. 22). We measured the binding affinity of 5 designs using surface plasmon resonance (SPR) and obtained affinity constants (K_D_s) between 150 and 790 nM (Fig. 6b-d), while knockout mutants and GLF scaffolds without the epitope did not exhibit binding to the ghrelin receptor ICL3-specific antibody (Fig. 6c-d).

**Fig. 6.**
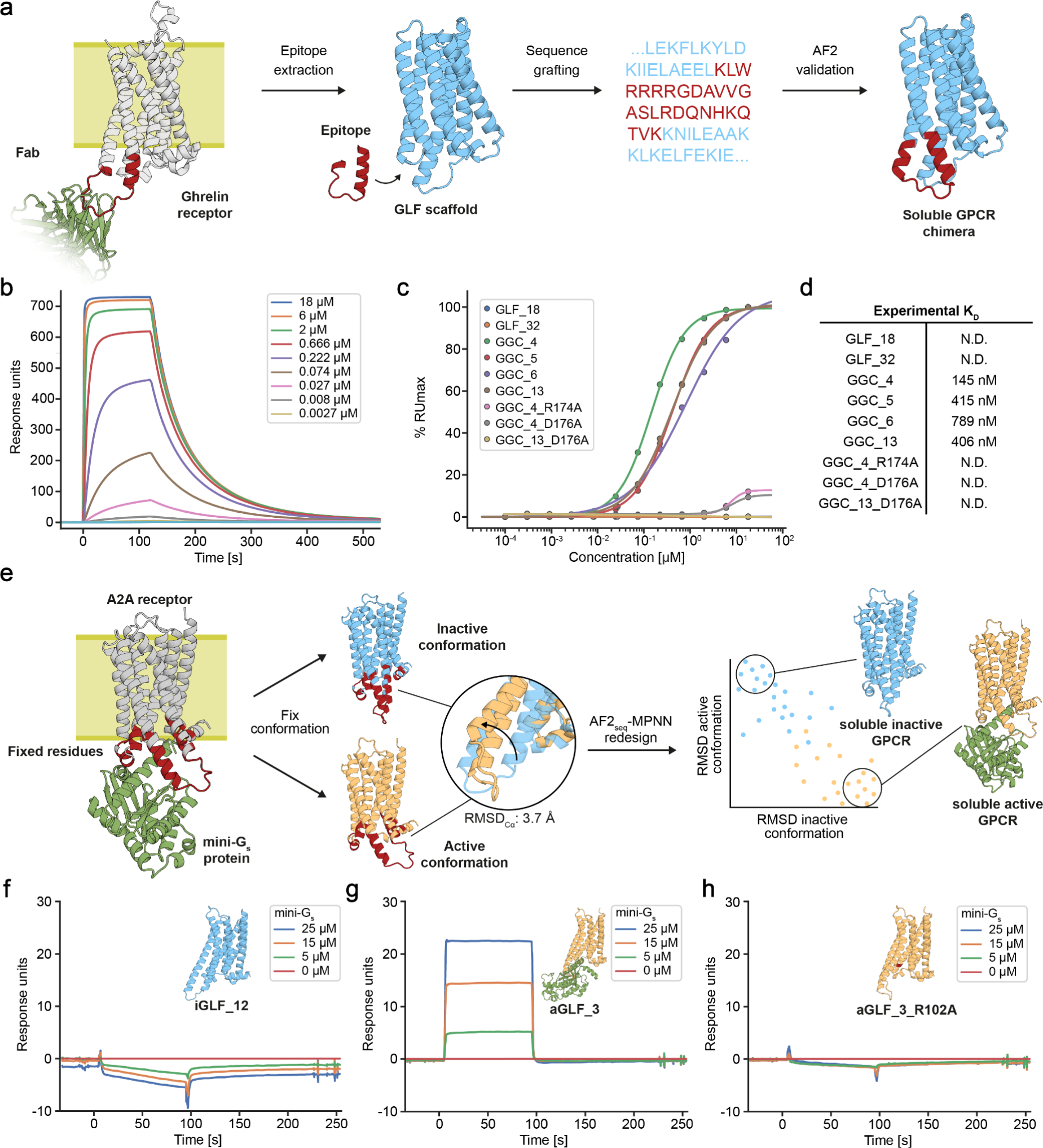
| Functionalization of soluble analogues of GPCR proteins. **a,** Design a workflow for the functionalization of soluble scaffolds via the grafting of the native epitope corresponding to the ICL3 loop of the Ghrelin GPCR receptor that can be probed with a Fab (PDB: 6KO5)^73^. **b,** Representative SPR sensorgram displaying the binding kinetics of increasing concentrations of Ghrelin targeting antibody binding to GGC_4. **c,** Binding affinities determined by SPR of the designed GGC constructs and corresponding negative controls. **d,** Table summarizing experimental affinity constants from data in panel c. **e,** Design a workflow for the conformation-specific design of the active^75^ and inactive^76^ form of the adenosine A2ACXCR2 receptor to facilitate or preclude mini-Gs protein binding. SPR sensorgram of the **f,** inactive form iGLF_12, **g,** active form aGLF_3, and **h,** binding knockout mutant of aGLF_3 soluble analogue.

An important part of GPCR receptor function is the activation of intracellular signaling pathways mediated by G-protein binding^57^. To recapitulate this functional aspect, we designed soluble analogues of the adenosine A2A receptor in a conformation-specific manner. This entailed the preservation of the G-protein binding site, including evolutionary conserved sequences, such as the DRY motif that is essential for receptor activation and G-protein binding^77^. This resulted in designs in both the active^75^ and inactive^76^ with identical fixed G-protein interacting residues (Fig. 6e, Methods). The active state is characterized by a shift of the transmembrane helix 6 (TM6) in an outwards rotational motion, thereby exposing the G-protein binding site^77^. We characterized 3 constitutively active (aGLF) and 3 constitutively inactive (iGLF) soluble GPCR analogues, and all were found to be monomeric and folded in solution (Extended Data Fig. 23). SPR binding experiments of miniG_s_-414^78^ revealed no binding to the iGLFs (Fig 6f, Extended Data Fig. 23), while a clear binding signal was observed for the aGLF designs (Fig 6g, Extended Data Fig. 23), however exact affinities could not be determined due to rapid interaction kinetics. To validate the specificity of the binding mode, we mutated the highly conserved DRY motif and observed complete abolishing of binding to the aGLFs (Fig 6h, Extended Data Fig. 23). Mutation of residues in the G-protein binding site, outside of the DRY motif, were also found to diminish or impair binding of miniGs-414 to aGLFs (Extended Data Fig. 23). These results indicate that specific functional states can be designed with high accuracy, while preserving critical evolutionarily conserved motifs within *de novo* designed scaffolds.

In summary, we present here a novel computational approach enabling conformation-specific design and functionalization of soluble membrane protein analogues through motif grafting and constrained design procedures, which could have a realm of important applications in the computational design of functional proteins.

## Conclusions

The robust computational design of novel protein folds remains a difficult endeavor. Here, we present a computational approach based on deep learning that through the generation of high-quality protein backbones enables the efficient search of non-natural sequences for a variety of protein topologies. The computational framework based on AF2_seq_-MPNN is flexible and generalizable, avoiding the need to perform fold-specific re-training or providing tedious parametric and symmetric design restraints for fold-conditioning. We designed and characterized several folds that have been very challenging to engineer with previous methods, achieving high experimental success rates in terms of soluble and folded designs. Structural characterization of the designs showed an exquisite accuracy of the computational models, both in terms of overall fold, as well as in the fine details of the side chain conformations, which are critical for the design of function. Additionally, we aimed to test the capability of the computational approach to provide the means to expand the soluble fold space and enable the design of analogues of protein topologies only found in membrane environments. By allowing full sequence design, we designed three different membrane fold analogs, including two with highly elaborate helical topologies (rhomboid protease and GPCR) and showed that such designs were folded and monomeric in solution. The experimental structures showed once again that the design method was very accurate, and that we recreate soluble analogues both for the rhomboid protease fold, as well as for the canonical 7 helical GPCR fold, which are not present in the soluble fold space. By doing so, we showed that membrane protein folds generally follow the same design principles as soluble protein folds, and that many of such folds can be readily designed in soluble form.

Moreover, we hypothesize that this could promote the designability of functional proteins by accessing a new plethora of folds that are not present in the soluble fold space. Another exciting perspective is the creation of soluble analogues of membrane proteins that retain many of the native features of the original membrane proteins, such as enzymatic or ligand-binding functions, which could greatly accelerate the study of their function in more biochemically accessible soluble formats. We demonstrated the potential of our method by incorporating native structural motifs into designed soluble analogues. By designing soluble analogues in the context of the natural functional site, we preserved even complex structural features of the sites, such as the extracellular beta-sheeted domains of claudins. Recent studies have identified claudins as potential targets for the treatment of certain types of cancers^79,80^, therefore the development of multiple classes of soluble claudins could potentially accelerate drug screening and serve as the basis for the design of claudin-based biologics. Specifically in GPCR drug development, it would be transformative to create soluble analogues in specific functional states that could be used for small molecule or antibody discovery campaigns. The precision of our design approach enabled conformational specific design for the active and inactive GPCR states, differentiated by subtle conformational changes. Consequently, our designs harbor identical G-protein binding sites, yet they uniquely either constitutively facilitate or preclude G-protein binding in solution. The computational design of specific conformational states that can mediate biological function remains an outstanding problem for which we provide a flexible and broadly applicable methodological workflow. Such an approach could constitute the basis for computational design strategies of proteins that are able to populate multiple conformational states in a predictable fashion, which is an important prerequisite to embedding complex functions in computationally designed proteins. From an applied perspective, the capability of creating membrane soluble analogues with native functional features could be critical to facilitate the development of novel drugs and therapies that target these challenging classes of proteins, which remain some of the most important drug targets. In summary, we present a deep learning approach for computational protein design that demonstrates the usefulness of high-quality structure representations to enable the effective exploration of new sequence spaces that can yield viable proteins and contribute to the expansion of the designable fold space that can ultimately impact our ability to design functional proteins.

## Materials and Methods

### AF2seq design protocol

#### Design target preparation

The design target structures were sourced from the Protein Data Bank (PDB) and included the following protein folds: Ig-like fold (3SD2), β-barrel fold (6D0T)^38^, TIM-barrel (5BVL)^50^, claudin (4P79)^55^, rhomboid protease (3ZEB)^56^, and GPCR (6FFI)^81^. Due to missing residue positions in the TIM-barrel, claudin, and GPCR X-ray structures, we used AlphaFold2 to predict the protein structure using the X-ray structure as a template. The claudin target contained a disordered region (residues 34 to 40) that we replaced with a three-glycine linker and the GPCR target contained a large disordered region (residues 875 to 896) which we replaced with a five glycine linker. Additionally, the GPCR X-ray target contained an endolysin domain that was inserted into one of the loops to enhance protein solubility for crystallization (residues 679 to 838). Therefore, we predicted the GPCR sequence without the endolysin domain using the X-ray target as a template.

#### Loss function

For the computation of error gradients, a composite loss function was used:

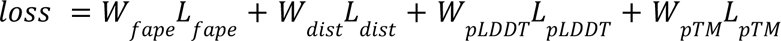

The loss function is represented as a combination of L, which denotes the value of the loss, and W, which denotes the weight of the loss. The Frame Aligned Point Error (FAPE) loss quantifies the L2 norm between the predicted C_α_ atoms and the target structure. The distogram (dist) loss is the cross entropy over the C_β_ distogram for non-glycine residues and the C_α_ distance in the case of glycine. The model confidence (pLDDT) loss of the C_α_ positions is computed by taking 1 – pLDDT, penalizing low confidence. And finally, the pTM-score (pTM) loss, which is a prediction confidence metric focused on global structural similarity. In this paper, the designs were generated using loss terms of W_fape_ = 1.0, W_pLDDT_ = 0.2 and W_pTM_ = 0.2. During initial trajectories, the W_dist_ was set to 0.5, while it was disabled during trajectory re-seeding (soft-starts, described below).

#### Gradient descent

As previously described by Goverde et al.^8^, the amino acid sequences were initialized based on the secondary structure of the target fold. The secondary structure assignments were then encoded in sequences using alanines for helix, valines for β-sheet and glycines for loop residues. This introduces a bias towards the correct local structure, aiding faster convergence of the design trajectories. In order to diversify the generated designs, 10% of the amino acids are randomly mutated in the initial sequence of each design trajectory. Subsequently, the sequence was passed through the AF2 networks, which generated five structures. These structures were then utilized to calculate the loss using the previously defined loss function. The error gradient was obtained by backpropagating the errors to the one-hot-encoded input, resulting in a 5×20×N error gradient, where N represents the sequence length. We then take the average of the five matrices to obtain the mean error gradient (20xN) which is used for gradient descent. A Position Specific Scoring Matrix (PSSM) of 20xN is updated using the ADAM optimizer^82^ with the normalized error gradient. Following the update, the PSSM undergoes a softmax function that transforms the matrix into a probability distribution of the amino acid identity for each position. The argmax function is subsequently used to determine the most probable amino acid identities per position, which are then utilized to construct the new input sequence for the next iteration. The cysteine residues in the PSSM are masked so the designed sequences do not contain any cysteines.

#### Model settings

AF2 was run in single sequence mode using the network configuration of the original AF2 ‘model_5_ptm’ for all five AF2 models with MSAs and templates disabled. For the design trajectories, we used zero recycles, meaning that each AF2 network was only executed once. For the claudin-1 and claudin-4 designs we only used models 1 and 2 with network configuration of the original AF2 ‘model_1_ptm’ with templates enabled. All design runs were executed on a single Nvidia Tesla V100 (32GB) GPU.

#### Computational design protocol

In each AF2-sequence design trajectory, 500 rounds of gradient descent optimization were performed (https://github.com/bene837/af2seq). Not all design trajectories of the claudin, rhomboid protease and GPCR converged. Hence, we sampled sequences from successful trajectories and introduced mutations, while disabling distogram loss. These sequences were then used as starting points for new design trajectories, we named soft-starts, which resulted in a higher convergence rate. All generated sequences were then predicted using AF2 with three recycles, followed by relaxation in an AMBER force-field^83,84^. This results in high quality structures which were used as an input for ProteinMPNN sequence generation. The total number of designs and number of designs passing *in silico* filtering is summarized in Table S2. For the design of the claudin-1 and claudin-4 functional analogues, we first predicted their structures using AF2 with MSAs and templates enabled, due to the lack of high-resolution experimental structures. The predictions were then used as structural templates for both design and reprediction, as the wild-type extracellular region could not be predicted by AF2 in single sequence mode. All sequence and sidechain information was removed from the template to reduce folding bias as much as possible. We tried several design strategies for the functional claudin design of which two were successful: 1) redesigning only the transmembrane surface, approximately 40% of the sequence; and 2) redesigning the entire transmembrane region, including core, approximately 60% of the sequence. The residue positions that were fixed can be found in the Extended data S1.

For conformation-specific design of GPCRs, we used the template of the adenosine A2A GPCR in the active conformation bound to mini-G_s_ (PDB: 5G53) and the inactive conformation (PDB: 3VGA) to design each state individually. We fixed residues interacting with the G-protein and the evolutionary conserved DRY motif during the design of each state, resulting in designs with an identical length and identical functional sites. For the design of the active conformation, we found that it was not possible to generate high confidence designs without the presence of G-protein, hence gradient descent and prediction was performed in the presence of the mini-G_s_ binder.

### Training of SolubleMPNN (MPNN_sol_)

The MPNN_sol_ model was trained on protein assemblies in the PDB (as of Aug 02, 2021) determined by X-ray crystallography or cryoEM to better than 3.5 Å resolution and with less than 10,000 residues. We followed training as described in the work by Daupauras et al.^15^ with the only modification of excluding annotated transmembrane PDB codes. The list of excluded PDBs and MPNN_sol_ model weights are available here: https://github.com/dauparas/ProteinMPNN/tree/main/soluble_model_weights.

### ProteinMPNN sequence redesign

The generated backbones by AF2_seq_ were used as input to proteinMPNN. For the vanilla proteinMPNN we used the provided model weights trained on a dataset with 0.1 Å gaussian noise^15^. For the biased proteinMPNN (referred in the main text as MPNN_bias_) we used a modified version of the script ‘submit_example_8.sh’ as provided on the proteinMPNN github mentioned above. We found the best results by giving a positive sampling bias to the polar amino-acids whilst giving a negative sampling bias to alanine. For soluble proteinMPNN (MPNN_sol_) we generated sequences with two different models which had different levels of noise during training (0.1 Å and 0.2 Å). For all proteinMPNN models we generated two sequences per AF2_seq_ designed backbone. No gaussian noise was added to the input backbone and cysteine residues were masked during the decoding process.

### Structural similarity calculations

The C_α_ atoms of the structures were aligned using the super imposer from the Biopython package^85^. The Root Mean Square Deviation of the C_α_ atoms (RMSD_Cα_) is calculated as the mean Euclidean distance between predicted and target C_α_ atom coordinates. The full atom RMSD (RMSD_fa_) is calculated by first aligning all atoms with the superimposer after which the mean Euclidean distance between atoms is computed. The Template Modeling (TM) scores were determined using TM-align^86^.

### Sequence diversity analysis

Sequence recovery is quantified as the number of positions at which the corresponding residue matches the residue in the target fold divided by the total number of residues in the sequence multiplied by 100%. The core residues are defined as residues with less than 20 Å^2^ Solvent-Accessible Surface Area (SASA) and surface residues are defined as residues with less than 20 Å^2^ SASA. The e-values were obtained through a protein BLAST search of the NCBI RefSeq database of 10/1/2022 with a maximum hit value of 1000.

### Surface hydrophobicity calculations

The fraction of surface hydrophobics were calculated using Rosetta^1^. First, all surface residues were identified using the layer selector; these residues are defined as residues with Solvent Accessible Surface Area (SASA) > 40 Å^2^. Of these surface residues we count the number of apolar amino acids (defined as “GPAVILMFYW”) and divide it by the total number of surface residues.

### Design filtering and selection

All generated sequences were predicted with AF2 using three recycles and a relaxation step in an AMBER force-field. Next the sequences were filtered using the following criteria: I) TM-score > 0.80 for all designs except the rhomboid protease. The rhomboid protease yielded slightly lower TM-scores in the design trajectory hence we chose a cutoff value 0.75 instead; II) pLDDT > 80 for all designs except the rhomboid protease (pLDDT > 75); III) an e-value threshold > 0.1 for sequence novelty.

### Structural fold similarity search

The fold similarity search was performed using FoldSeek^87^ on the SCOP-database^21^ (downloaded March 2023). For each of the design target folds, an exhaustive search based on TM-score alignment was performed. The SCOP-database contains globular and membrane domain annotations which were used for the hit classification.

### Fold complexity calculations

Relative Contact Order (RCO) is calculated at the secondary structure level by computing the residue distance in the sequence between secondary structures for all pairs within 8 Å of each other, and then averaging these distances for all contacts that are more than 4 residues apart. To ensure consistency in secondary structure annotations across all structures, we employed DSSP for the determination of secondary structural elements^21^. The de novo protein dataset comprised 70 helical proteins, 6 beta sheet proteins, and 42 proteins containing both alpha helices and beta sheets^43,88^. Meanwhile, the natural protein dataset consisted of 1,000 proteins, randomly selected from the entire collection of proteins in the CATH dataset (version 4.3)^89^.

### Molecular dynamics-based backbone perturbation

All molecular dynamics (MD) simulations were performed using the Groningen Machine for Chemical Simulations (GROMACS) 2021.4 software package^90^. For the unbiased simulations in water, the Amber ff99SB-ILDN force field for proteins^91^ was used. The simulations were carried out under *NPT* conditions with a leapfrog integration scheme and a time step of 2.0 fs. Rhombic dodecahedron (triclinic) periodic boundary conditions were applied and the TIP3P^91^ water model was used as solvent. Temperature coupling using stochastic velocity rescaling to two separate temperature baths for the protein and for the water solvent was applied with a reference temperature of 300 K and a relaxation time of 0.1 ps. The pressure was coupled isotropically to a Parrinello-Rahman barostat at 1.0 bar with a coupling constant of 2.0 ps and an isothermal compressibility of 0.45 nm^2^ N^-1^. For both the short-range electrostatic- and van der Waals interactions, a single cutoff distance of 0.9 nm was used. The long-range electrostatics were calculated by the particle mesh Ewald (PME) algorithm with a Fourier spacing of 0.12 nm. The linear constraint solver (LINCS) algorithm was used to impose constraints on the bond lengths with fourth order expansion. Preceding the simulations, the solvated protein structures were energy minimized with a steepest descent algorithm, until the maximum force was below 100 kJ mol^-1^ nm^-1^. For the unbiased simulations in the POPC lipid bilayer (a mimic for a cellular membrane), the GROMOS 54A8 force field was used in combination with lipid parameters from the 1-Palmitoyl-2-oleoylphosphatidylcholine (POPC) model of Marzuoli *et al.*^92^. The simulations were carried out under *NPT* conditions with a leapfrog integration scheme and a time step of 2 fs. Rectangular periodic boundary conditions were applied and the SPC water model was used as solvent. Nose-Hoover temperature coupling was applied to two separate temperature baths at 323 K, one for the protein and the POPC bilayer and one for the solvent, with a relaxation time of 0.5 ps. The pressure was coupled semi-isotropically to a Parrinello-Rahman barostat at 1.0 bar with a coupling constant of 2.0 ps and an isothermal compressibility of 0.45 nm^2^ N^-1^ was applied. For both the short-range electrostatic and van der Waals interactions, a single cutoff of 1.2 nm was used. The long-range electrostatics were treated using the PME algorithm with a Fourier spacing of 0.16 nm. The bond lengths were constrained with the linear constraint solver (LINCS) algorithm. The simulation systems contained 512 POPC molecules in a bilayer (256 per leaflet). These lipids were packed around the embedded protein structures in the center of the simulation box (2.4963 nm x 13.0172 nm x 13.7189) using the InflateGRO methodology [KANDT2007475], as described by Lemkul^93^. Center-of-mass (COM) motion removal was applied in every simulation step to remove the motion of the bilayer and protein relative to the solvent. Preceding the simulations, the solvated simulation systems were energy minimized with a steepest descent algorithm, until the maximum force was below 1000 kJ mol^-1^ nm^-1^. For all simulations, the energy minimization was followed by 100 ps *NVT* thermalisation and 1.0 ns *NPT* equilibration. After equilibration, initial velocities were generated using a random number generator. Three unbiased MD runs were performed for 11 ns each and the first 1.0 ns of each simulation was discarded, resulting in 30 ns of sampling for each starting structure. Trajectory frames were extracted every 100 ps.

### Design solubility and binding screen

Designs were synthesized as gene fragments by Twist Bioscience and Gibson cloned between the NheI and XhoI sites in pET21b vector with C-terminal His_6_-tag. Designs were transformed into *E. coli* BL21 DE3 cells and expressed overnight with 1 mM IPTG at 18 °C in 1 ml LB medium supplemented with 100 μg/ml ampicillin in a 96-well format. After expression, cells were chemically lysed, cell debris was pelleted, and the supernatant was incubated with 50 μl of equilibrated Ni-NTA agarose (QIAGEN) beads. After incubation, unbound supernatant is discarded, beads were washed 5 times with 50 mM Tris-HCl pH 7.5, 500 mM KCl, 15 mM imidazole, and designs were eluted using buffer containing 50 mM Tris-HCl pH 7.5, 500 mM KCl, 400 mM imidazole. Elutions were analyzed using SDS-PAGE and designs were denoted soluble if the appropriate protein band is clearly visible following Coomassie staining. For binding screens, Pierce™ Protein A/G Agarose (Thermo Scientific) beads were used instead, and imidazole is omitted from the washing step. Ten µg of Fab were added to each lysate, allowing to pull-down interacting constructs. Complexes were eluted with 0.1 M glycine pH 2.5 and analyzed using SDS-PAGE.

### Transplantation of natural epitopes onto soluble scaffolds

Compatible epitopes were identified via a Foldseek search^1^ of the PDB database, using soluble scaffolds as queries. Hits with TM-scores above 0.7 and high structural similarity around the desired epitope were then superimposed in a structure visualization software such as PyMOL or ChimeraX. Varying lengths of the epitope were selected for transplantation, encompassing either only interaction sites, entire loops, or overlapping parts of the supporting secondary structures. The sequence of the overlaid epitope was then pasted into the overlapping region of interest within the soluble scaffold. The resulting chimeric sequences were predicted using AF2 in single sequence mode. Structures with high pLDDT (>90) and high TM-scores to the starting scaffold were manually inspected to verify the placement of the epitope. Finally, a subset of constructs in different soluble scaffolds were selected for experimental testing.

### Surface Plasmon Resonance (SPR) binding assay

SPR measurements were carried out on the Biacore 8K system (Cytiva) in HBS-EP+ buffer (10 mM HEPES pH 7.4, 150 mM NaCl, 3 mM EDTA, 0.005% (v/v) Surfactant P20 Cytiva). The antibody at a concentration of 5 µg/ml was immobilized on a CM5 sensor chip (Cytiva) via amide coupling in 10 mM NaOAc pH 4.5 for 250s at a flow rate of 10 µl/min (700-1500 response units immobilized). Purified miniGs was immobilized with a contact time of 200s (final immobilization level of 300 response units). Serial dilutions of the designed chimeras at concentrations ranging from 18 µM to 0.1 nM, and 0 nM were sequentially injected and the immobilized antibody was regenerated with 10 mM Glycine-HCl pH 2.5 for 30s at a flow rate of 30 µl/min in between injections. GPCRs designed for the active or inactive state were injected at concentrations of 0, 5, 15, and 25 µM for 90s at 30 µl/min followed by a dissociation phase of 120s at identical flow rate. The immobilized miniGs was not regenerated in between cycles. The analytes were allowed to bind to the immobilized antibody for 120s at a flow rate of 30 µl/min followed by a dissociation phase lasting 400s at identical flow rate. Binding curves were fitted with a 1:1 Langmuir binding model in the Biacore 8K analysis software. Steady-state response units were plotted against analyte concentration and a sigmoid function was fitted to the experimental data in Python 3.9 to derive the K_D_.

### Bio-Layer Interferometry (BLI)

For BLI studies of claudins we used synthetic claudin-_His_ and tagless *Clostridium perfringens* enterotoxin (CpE) both equilibrated in a buffer consisting of 20 mM Tris pH 7.4, 100 mM NaCl, and 5% glycerol. BLI was performed at 25°C in 96-well black flat bottom plates (Greiner) using an acquisition rate of 5 Hz averaged by 20 using an Octet© R8 System (FortéBio/Sartorius), with assays designed and setup using Blitz Pro 1.3 Software. Binding experiments consisted of the following steps: sensor equilibration (30 seconds), loading (300 seconds), baseline (180 seconds), and association and dissociation (120-300 seconds each). Experiments were conducted by immobilizing 1.5-3 µM of synthetic claudin-_His_ on NiNTA (Dip and Read) sensors and quantifying their binding to 0.05-5 µM CpE. Association and dissociation times for the two claudin-1 designs were performed for 120 seconds as they exhibited rapid on and off rates, while for the claudin-4 design these times were extended to 300 seconds to capture the slower off rates. Data were fit to a 1:1 binding model using the Octet^®^ Analysis Studio (Sartorius), which generated the K_D_ from k_on_ and k_off_ rates. At the protein concentrations used, no significant non-specific binding of CpE to NiNTA sensors were detected.

### Protein expression, purification and characterization

Designed proteins were expressed in *E. coli* BL21 (DE3) (Novagen) for 16 h at 18 °C. Bacterial pellets were resuspended and sonicated in 50 mM Tris-HCl pH 7.5, 500 mM KCl, 15 mM imidazole buffer supplemented with 1 mg/ml lysozyme, 1 mM PMSF, and 1 μg/ml DNase. Cell lysates were clarified using ultracentrifugation and loaded on a 10 ml Ni-NTA Superflow column (QIAGEN) and washed with 7 column volumes of 50 mM Tris-HCl pH 7.5, 500 mM NaCl, 10 mM imidazole. Designs were eluted with 10 column volumes of 50 mM Tris-HCl pH 7.5, 500 mM NaCl, 500 mM imidazole. Main protein fractions were concentrated and injected onto a Superdex 75 16/600 gel filtration column (GE Healthcare) in PBS buffer. CLN4_20 for cryoEM studies was purified using a S200 10/300 GL column (GE Healthcare) and eluted in 20 mM HEPES pH 8.0, 150 mM NaCl. Protein fractions were concentrated, flash frozen in liquid nitrogen and stored at −80 °C. Molar mass and homogeneity were confirmed using SEC-MALS. Folding, secondary structure content, and melting temperatures were assessed using circular dichroism in a Chirascan V100 instrument from Applied Photophysics. miniGs construct 414 was expressed and purified as described previously^78^.

### Antibody and Fab expression and purification

IgG antibodies and Fabs were expressed in 25 ml cultures of Expi293 cells with Invitrogen ExpiFectamine™ 293 Transfection Kit (A14525) following supplier’s recommendations. After 6 days of secretion, the cell culture supernatant was collected, loaded on a 5 ml Ni-NTA Superflow column (QIAGEN), and eluted in 50 mM Tris-HCl pH 7.5, 500 mM NaCl, 500 mM imidazole buffer. The eluate was further purified using a Superdex 200 16/600 gel filtration column in PBS. The protein eluted as a single peak at expected retention volume. Collected fractions were concentrated to 1 mg/ml, snap-frozen in liquid nitrogen, and stored at −80 °C.

### Protein crystallization and structure determination

The TIM barrel TBF_24 design was crystallized in the P2_1_2_1_2_1_ space group using the sitting drop vapor diffusion setup at 4 °C in 0.1 M Na_3_ Citrate pH 4.0, 1 M LiCl, 20% PEG 6000 buffer. The Claudin-like CLF_4 design was crystallized in the P1 space group using the sitting drop vapor diffusion method at 4 °C in 0.1 M Na_3_ Citrate pH 5.0, 0.1 M Na/K phosphate pH 5.5, 0.1 M RbCl, 25% v/v PEG smear medium (BCS Screen, Molecular Dimensions). The Rhomboid protease-like RPF_9 design was crystallized in the P2_1_2_1_2_1_ space group using the sitting drop vapor diffusion setup at 4 °C in 0.1 M HEPES pH 7.8, 0.15 M Na_3_ Citrate dihydrate, 25% v/v PEG smear low (BCS Screen, Molecular Dimensions). The GPCR-like GLF_18 design was crystallized in the P1 space group using the sitting drop vapor diffusion setup at 4 °C in Na Phosphate-Citrate pH 4.2, 0.2 M LiSO4, 20% PEG 1000 buffer. The GPCR-like GLF_32 design was crystallized in the P2_1_2_1_2_1_ space group using the sitting drop vapor diffusion setup at 4°C in 0.1 M Na Acetate pH 5.5, 0.2 M KBr, 25% PEG MME 2000 buffer. Crystals were cryoprotected in the crystallization buffer supplemented with 20% glycerol and flash-cooled in liquid nitrogen. Diffraction data was measured at the beamline PXI (X06SA) of the Swiss Light Source (Paul Scherrer Institute, Villigen, Switzerland) and the MASSIF-1 beamline of the The European Synchrotron Radiation Facility, Grenoble, France at a temperature of 100 K. Data was processed using the autoPROC package^94^, and phases were obtained by molecular replacement using the Phaser module of the Phenix package^95^. Atomic model adjustment and refinement was completed using COOT^94^ and Phenix.refine^96^. The quality of refined models was assessed using MolProbity^97^. Structural figures were generated using PyMOL (Schrödinger, LLC; https://www.pymol.org/) and ChimeraX.

### cryoEM structure determination of CLN4-20 in complex with cCpE

Expression and purification of cCpE, COP-2 Fab, and the anti-Fab nanobody was performed as described previously^72^. Concentrated CLN4_20 was first complexed with cCpE followed by COP-2 in a 1:1.2:1 molar excess. Next, the anti-Fab nanobody was added at 1.3 molar excess of COP-2 and incubated on ice for 30 min, concentrated using a 50 kDa MWCO concentrator (Millipore), and subjected to SEC using a Superdex 200 increase 10/300 GL column (GE Healthcare) pre-equilibrated in 20 mM HEPES pH 8.0, 150 mM NaCl. The purified protein complex was concentrated to 5 mg/mL using a 50 kDa MWCO concentrator (Millipore) and filtered with 0.2 µm filter.

UltraAuFoil 1.2/1.3 grids (Quantifoil) were glow discharged for 30 s at 15 mA in a Pelco easiGlow (Ted Pella Inc) instrument and vitrified using Leica GP2 instrument (Leica microsystems). 3.5 µL of the complex was applied onto grids and blotted for 3 s at 4°C under 100% humidity, then plunge frozen into liquid ethane. Grid screening and data collection was performed on 200 kV Glacios 2 Cryo-TEM (ThermoFisher Scientific) with Falcon 4i Direct electron detector and EPU software (ThermoFisher Scientific) at Hauptman-Woodward Medical Research Institute. Total 1159 movies were collected at a physical pixel size of 0.884 Å, electron dose of 49.4 e/Å^2^ fractioned over 93 frames.

Movies were processed, patch motion corrected, and patch CTF estimated in cryoSPARC. Blob picking generated a suitable template for an initial 3D volume which was used to produce 2D projections for template picking, followed by 2D classification, *ab initio* reconstruction, and 3D refinement, which resulted in a cryoEM density resolved to a resolution of 4.1 Å. Structural coordinates for the complex between CLN4_20, cCpE, and COP-2 Fab from PDB ID: 7TDM^72^ were rigid body docked into the 4.1 Å map. The nanobody from PDB ID: 8U4V was docked onto the L chain of COP-2 to complete the four-protein complex. Once the model was manually fitted, each protein chain was real-space refined in Coot to place the structure within the cryo-EM map volume. Final model refinement was conducted using Namdinator^98^, followed by real-space refinement using Phenix phenix.real_space_refine^94^. Table S2 shows data collection, refinement, and validation statistics for the CLN-4_20/cCpE/COP-2/Nb structure.

## Acknowledgments

We thank SCITAS at EPFL for support in running design trajectories. We thank Beat Blattmann (Protein Crystallization Center, UZH, Zurich, Switzerland) for performing initial crystallization screens. We thank Florence Pojer, Kelvin Lau, and Amédé Larabi (Protein Production and Structure Characterization Core Facility, EPFL, Switzerland) for help with crystal optimization and data processing. We thank Takashi Tomizaki (Swiss Light Source, X06SA beamline, Paul Scherrer Institute, Villigen, Switzerland), and Didier Nurizzo (The European Synchrotron Radiation Facility, MASSIF-1 beamline, Grenoble, France) for assistance with crystallographic data collection. We thank Chris Tate (MRC Laboratory of Molecular Biology, Cambridge) for kindly donating the expression plasmid of miniGs-414.

## Funding

MP was supported by the Peter und Traudl Engelhorn Stiftung. BEC was supported by the Swiss National Science Foundation, the NCCR in Chemical Biology, the NCCR in Molecular Systems Engineering, and the ERC Starting grant no. 716058. JD was supported by a gift from Microsoft and DB was supported by the Howard Hughes Medical Institute. SO and SK were supported by NIH DP5OD026389, NSF MCB2032259, and Amgen. AJV was supported by the National Institute of General Medical Sciences of the NIH under Award Number R35GM138368.

## Author contributions

Conceptualization: CAG, MP, BEC

Methodology: CAG, MP, SK, JD, DB, SO, BEC

Investigation: CAG, MP, NG, PEMB, LJD, SG, SR, SK, JC, CS, SK, JD

Visualization: CAG, MP, NG, LJD, JD, BEC

Funding acquisition: DB, AJV, SO, BEC

Project administration: CAG, MP, BEC

Supervision: DB, AJV, SO, BEC

Writing – original draft: CAG, MP, LJD, BEC

Writing – review & editing: CAG, MP, NG, LJD, SK, JD, DB, SO, BEC

## Competing interests

Authors declare that they have no competing interests.

## Data and materials availability

All data are available in the main text or as supplementary materials. Atomic coordinates and structure factors of the reported X-ray structures have been deposited in the Protein Data Bank under accession numbers 8OYS (TBF_24), 8OYV (CLF_4), 8OYW (RPF_9), 8OYX (GLF_18), and 8OYY (GLF_32). Af2seq code is available at https://github.com/bene837/af2seq. AlphaFold2 model weights used for design and predictions can be downloaded from https://storage.googleapis.com/alphafold/alphafold_params_2021-07-14.tar. ProteinMPNN, along with soluble trained weights is available at https://github.com/dauparas/ProteinMPNN.

## Notes

### Competing Interest Statement

The authors have declared no competing interest.

### Summary of Updates

Functionalization of the soluble analogs has been added to the manuscript.

## References

1. Leman, J. K. et al. Macromolecular modeling and design in Rosetta: recent methods and frameworks. Nat Methods 17, 665–680 (2020).

2. Jumper, J. et al. Highly accurate protein structure prediction with AlphaFold. Nature 596, 583–589 (2021).

3. Baek, M. et al. Accurate prediction of protein structures and interactions using a 3-track neural network. Sci New York N Y 373, 871–876 (2021).

4. Huang, B. et al. A backbone-centred energy function of neural networks for protein design. Nature 602, 523–528 (2022).

5. Lin, Y. & AlQuraishi, M. Generating Novel, Designable, and Diverse Protein Structures by Equivariantly Diffusing Oriented Residue Clouds. (2023).

6. Frank, C., et al. Efficient and scalable de novo protein design using a relaxed sequence space. bioRxiv (2023) doi:10.1101/2023.02.24.529906.

7. Anishchenko, I. et al. De novo protein design by deep network hallucination. Nature 600, 547–552 (2021).

8. Goverde, C., Wolf, B., Khakzad, H., Rosset, S. & Correia, B. E. De novo protein design by inversion of the AlphaFold structure prediction network. Protein Sci e4653 (2023) doi:10.1002/pro.4653.

9. Wicky, B. I. M. et al. Hallucinating symmetric protein assemblies. Science 0, eadd1964 (2022).

10. Watson, J. L. et al. De novo design of protein structure and function with RFdiffusion. Nature 1–3 (2023) doi:10.1038/s41586-023-06415-8.

11. Wang, J. et al. Scaffolding protein functional sites using deep learning. Science 377, 387–394 (2022).

12. Gainza, P. et al. De novo design of protein interactions with learned surface fingerprints. Nature 617, 1–9 (2023).

13. Ingraham, J., Garg, V., Barzilay, R. & Jaakkola, T. Generative models for graph-based protein design. Advances in Neural Information Processing Systems 32 (NeurIPS 2019) 32, 15741–15752 (2019).

14. Anand, N. et al. Protein sequence design with a learned potential. Nat Commun 13, 746 (2022).

15. Dauparas, J. et al. Robust deep learning–based protein sequence design using ProteinMPNN. Science 0, eadd2187 (2022).

16. Yim, J., et al. SE(3) diffusion model with application to protein backbone generation. Arxiv (2023) doi:10.48550/arxiv.2302.02277.

17. Wu, K. E., et al. Protein structure generation via folding diffusion. (2022).

18. Trippe, B. L., et al. Diffusion probabilistic modeling of protein backbones in 3D for the motif-scaffolding problem. arXiv (2022).

19. Anand, N. & Achim, T. Protein Structure and Sequence Generation with Equivariant Denoising Diffusion Probabilistic Models. arXiv (2022).

20. Norn, C. et al. Protein sequence design by conformational landscape optimization. P Natl Acad Sci Usa 118, e2017228118 (2021).

21. Andreeva, A., Kulesha, E., Gough, J. & Murzin, A. G. The SCOP database in 2020: expanded classification of representative family and superfamily domains of known protein structures. Nucleic Acids Res 48, D376–D382 (2019).

22. Slovic, A. M., Kono, H., Lear, J. D., Saven, J. G. & DeGrado, W. F. Computational design of water-soluble analogues of the potassium channel KcsA. Proc National Acad Sci 101, 1828–1833 (2004).

23. Qing, R. et al. Protein Design: From the Aspect of Water Solubility and Stability. Chem. Rev. 122, 14085–14179 (2022).

24. Slovic, A. M., Lear, J. D. & DeGrado, W. F. De novo design of a pentameric coiled-coil: decoding the motif for tetramer versus pentamer formation in water-soluble phospholamban*. J. Pept. Res. 65, 312–321 (2005).

25. Roosild, T. P. & Choe, S. Redesigning an integral membrane K+ channel into a soluble protein. Protein Eng. Des. Sel. 18, 79–84 (2005).

26. Zhang, S. et al. QTY code enables design of detergent-free chemokine receptors that retain ligand-binding activities. Proc. Natl. Acad. Sci. 115, E8652–E8659 (2018).

27. Moffat, L., Greener, J. G. & Jones, D. T. Using AlphaFold for Rapid and Accurate Fixed Backbone Protein Design. Biorxiv 2021.08.24.457549 (2021) doi:10.1101/2021.08.24.457549.

28. Jendrusch, M., Korbel, J. O. & Sadiq, S. K. AlphaDesign: A de novo protein design framework based on AlphaFold. Biorxiv 2021.10.11.463937 (2021) doi:10.1101/2021.10.11.463937.

29. Smith, C. A. & Kortemme, T. Backrub-Like Backbone Simulation Recapitulates Natural Protein Conformational Variability and Improves Mutant Side-Chain Prediction. J Mol Biol 380, 742–756 (2008).

30. Woof, J. M. & Burton, D. R. Human antibody–Fc receptor interactions illuminated by crystal structures. Nat Rev Immunol 4, 89–99 (2004).

31. Williams, A. F. & Barclay, A. N. The Immunoglobulin Superfamily—Domains for *Cell* Surface Recognition. Annu Rev Immunol 6, 381–405 (1988).

32. Jost, C. & Plückthun, A. Engineered proteins with desired specificity: DARPins, other alternative scaffolds and bispecific IgGs. Curr Opin Struc Biol 27, 102–112 (2014).

33. Hecht, M. H. De novo design of beta-sheet proteins. Proc National Acad Sci 91, 8729–8730 (1994).

34. Bork, P., Holm, L. & Sander, C. The Immunoglobulin Fold Structural Classification, Sequence Patterns and Common Core. J Mol Biol 242, 309–320 (1994).

35. Marcos, E. et al. De novo design of a non-local β-sheet protein with high stability and accuracy. Nat Struct Mol Biol 25, 1028–1034 (2018).

36. Chidyausiku, T. M. et al. De novo design of immunoglobulin-like domains. Nat Commun 13, 5661 (2022).

37. Weber, B. et al. A single residue switch reveals principles of antibody domain integrity. J Biol Chem 293, 17107–17118 (2018).

38. Dou, J. et al. De novo design of a fluorescence-activating β-barrel. Nature 561, 485–491 (2018).

39. Vorobieva, A. A. et al. De novo design of transmembrane β barrels. Science 371, (2021).

40. Kipnis, Y. et al. Design and optimization of enzymatic activity in a de novo β-barrel scaffold. Protein Sci 31, (2022).

41. Tinberg, C. E. et al. Computational design of ligand-binding proteins with high affinity and selectivity. Nature 501, 212–216 (2013).

42. Bick, M. J. et al. Computational design of environmental sensors for the potent opioid fentanyl. Elife 6, e28909 (2017).

43. Verkuil, R. et al. Language models generalize beyond natural proteins. bioRxiv 2022.12.21.521521 (2022) doi:10.1101/2022.12.21.521521.

44. Gerlt, J. A. New wine from old barrels. Nat Struct Biol 7, 171–173 (2000).

45. Sterner, R. & Höcker, B. Catalytic Versatility, Stability, and Evolution of the (βα)8-Barrel Enzyme Fold. Chem Rev 105, 4038–4055 (2005).

46. Röthlisberger, D. et al. Kemp elimination catalysts by computational enzyme design. Nature 453, 190–195 (2008).

47. Romero-Romero, S., Kordes, S., Michel, F. & Höcker, B. Evolution, folding, and design of TIM barrels and related proteins. Curr Opin Struc Biol 68, 94–104 (2021).

48. Houbrechts, A. et al. Second-generation octarellins: two new de novo (beta/alpha)8 polypeptides designed for investigating the influence of beta-residue packing on the alpha/beta-barrel structure stability. Protein Eng 8, 249–59 (1995).

49. Nagarajan, D., Deka, G. & Rao, M. Design of symmetric TIM barrel proteins from first principles. Bmc Biochem 16, 18 (2015).

50. Huang, P.-S. et al. De novo design of a four-fold symmetric TIM-barrel protein with atomic-level accuracy. Nat Chem Biol 12, 29–34 (2016).

51. Chu, A. E., Fernandez, D., Liu, J., Eguchi, R. R. & Huang, P.-S. De Novo Design of a Highly Stable Ovoid TIM Barrel: Unlocking Pocket Shape towards Functional Design. Biodesign Res 2022, 1–13 (2022).

52. Mitra, K., Steitz, T. A. & Engelman, D. M. Rational design of ‘water-soluble’ bacteriorhodopsin variants. Protein Eng Des Sel 15, 485–492 (2002).

53. Perez-Aguilar, J. M. et al. A Computationally Designed Water-Soluble Variant of a G-Protein-Coupled Receptor: The Human Mu Opioid Receptor. Plos One 8, e66009 (2013).

54. Li, H., Cocco, M. J., Steitz, T. A. & Engelman, D. M. Conversion of Phospholamban into a Soluble Pentameric Helical Bundle †. Biochemistry-us 40, 6636–6645 (2001).

55. Suzuki, H. et al. Crystal Structure of a Claudin Provides Insight into the Architecture of Tight Junctions. Science 344, 304–307 (2014).

56. Tichá, A., Collis, B. & Strisovsky, K. The Rhomboid Superfamily: Structural Mechanisms and Chemical Biology Opportunities. Trends Biochem Sci 43, 726–739 (2018).

57. Katritch, V., Cherezov, V. & Stevens, R. C. Structure-Function of the G Protein–Coupled Receptor Superfamily. Annu Rev Pharmacol 53, 531–556 (2013).

58. Suzuki, H., Tani, K. & Fujiyoshi, Y. Crystal structures of claudins: insights into their intermolecular interactions. Ann Ny Acad Sci 1397, 25–34 (2017).

59. Li, J. Targeting claudins in cancer: diagnosis, prognosis and therapy. Am J Cancer Res 11, 3406–3424 (2021).

60. Vonrhein, C. et al. Data processing and analysis with the autoPROC toolbox. Acta Crystallogr Sect D Biological Crystallogr 67, 293–302 (2011).

61. Wang, Y., Zhang, Y. & Ha, Y. Crystal structure of a rhomboid family intramembrane protease. Nature 444, 179–180 (2006).

62. Urban, S., Lee, J. R. & Freeman, M. Drosophila Rhomboid-1 Defines a Family of Putative Intramembrane Serine Proteases. Cell 107, 173–182 (2001).

63. Hauser, A. S. et al. Pharmacogenomics of GPCR Drug Targets. Cell 172, 41–54.e19 (2018).

64. Hauser, A. S., Attwood, M. M., Rask-Andersen, M., Schiöth, H. B. & Gloriam, D. E. Trends in GPCR drug discovery: new agents, targets and indications. Nat Rev Drug Discov 16, 829–842 (2017).

65. Laguerre, M., Saux, M., Dubost, J. P. & Carpy, A. Mlpp: A Program for the Calculation of Molecular Lipophilicity Potential in Proteins. Pharm Pharmacol Commun 3, 217–222 (1997).

66. Rovati, G. E., Capra, V. & Neubig, R. R. The Highly Conserved DRY Motif of Class A G Protein-Coupled Receptors: Beyond the Ground State. Mol Pharmacol 71, 959–964 (2007).

67. Konvicka, K., Guarnieri, F., Ballesteros, J. A. & Weinstein, H. A Proposed Structure for Transmembrane Segment 7 of G Protein-Coupled Receptors Incorporating an Asn-Pro/Asp-Pro Motif. Biophys J 75, 601–611 (1998).

68. Srinivasan, M. & Dunker, A. K. Proline Rich Motifs as Drug Targets in Immune Mediated Disorders. Int J Peptides 2012, 634769 (2012).

69. Vecchio, A. J., Rathnayake, S. S. & Stroud, R. M. Structural basis for Clostridium perfringens enterotoxin targeting of claudins at tight junctions in mammalian gut. Proc. Natl. Acad. Sci. United States Am. 118, e2024651118 (2021).

70. Saitoh, Y. et al. Structural insight into tight junction disassembly by Clostridium perfringens enterotoxin. Science 347, 775–778 (2015).

71. Sonoda, N. et al. Clostridium perfringens Enterotoxin Fragment Removes Specific Claudins from Tight Junction Strands. J. Cell Biol. 147, 195–204 (1999).

72. Orlando, B. J. et al. Development, structure, and mechanism of synthetic antibodies that target claudin and Clostridium perfringens enterotoxin complexes. J. Biol. Chem. 298, 102357 (2022).

73. Shiimura, Y. et al. Structure of an antagonist-bound ghrelin receptor reveals possible ghrelin recognition mode. Nat. Commun. 11, 4160 (2020).

74. Liu, H. et al. Structural basis of human ghrelin receptor signaling by ghrelin and the synthetic agonist ibutamoren. Nat. Commun. 12, 6410 (2021).

75. Carpenter, B., Nehmé, R., Warne, T., Leslie, A. G. W. & Tate, C. G. Structure of the adenosine A2A receptor bound to an engineered G protein. Nature 536, 104–107 (2016).

76. Hino, T. et al. G-protein-coupled receptor inactivation by an allosteric inverse-agonist antibody. Nature 482, 237–240 (2012).

77. Hauser, A. S. et al. GPCR activation mechanisms across classes and macro/microscales. Nat. Struct. Mol. Biol. 28, 879–888 (2021).

78. Carpenter, B. & Tate, C. G. Engineering a minimal G protein to facilitate crystallisation of G protein-coupled receptors in their active conformation. *Protein Eng.*, Des. Sel. 29, 583–594 (2016).

79. Cherradi, S. et al. Antibody targeting of claudin-1 as a potential colorectal cancer therapy. J. Exp. Clin. Cancer Res. : CR 36, 89 (2017).

80. Fatima, I. et al. Identification and characterization of a first-generation inhibitor of claudin-1 in colon cancer progression and metastasis. Biomed. Pharmacother. 159, 114255 (2023).

81. Christopher, J. A. et al. Structure-Based Optimization Strategies for G Protein-Coupled Receptor (GPCR) Allosteric Modulators: A Case Study from Analyses of New Metabotropic Glutamate Receptor 5 (mGlu5) X-ray Structures. J Med Chem 62, 207–222 (2019).

82. Kingma, D. P. & Ba, J. Adam: A Method for Stochastic Optimization. Arxiv (2014) doi:10.48550/arxiv.1412.6980.

83. Hornak, V. et al. Comparison of multiple Amber force fields and development of improved protein backbone parameters. *Proteins: Struct., Funct.*, Bioinform. 65, 712–725 (2006).

84. Case, D. A. et al. The Amber biomolecular simulation programs. J. Comput. Chem. 26, 1668–1688 (2005).

85. Cock, P. J. A. et al. Biopython: freely available Python tools for computational molecular biology and bioinformatics. Bioinformatics 25, 1422–1423 (2009).

86. Zhang, Y. & Skolnick, J. TM-align: a protein structure alignment algorithm based on the TM-score. Nucleic Acids Res 33, 2302–2309 (2005).

87. Kempen, M. van, et al. Fast and accurate protein structure search with Foldseek. Nat Biotechnol 1–4 (2023) doi:10.1038/s41587-023-01773-0.

88. Woolfson, D. N. A Brief History of De Novo Protein Design: Minimal, Rational, and Computational. J Mol Biol 433, 167160 (2021).

89. Sillitoe, I. et al. CATH: increased structural coverage of functional space. Nucleic Acids Res 49, D266–D273 (2020).

90. Lindahl, E., Hess, B. & Spoel, D. van der. GROMACS 3.0: a package for molecular simulation and trajectory analysis. J Mol Model 7, 306–317 (2001).

91. Lindorff-Larsen, K. et al. Improved side-chain torsion potentials for the Amber ff99SB protein force field: Improved Protein Side-Chain Potentials. Proteins Struct Funct Bioinform 78, 1950–1958 (2010).

92. Marzuoli, I., Margreitter, C. & Fraternali, F. Lipid Head Group Parameterization for GROMOS 54A8: A Consistent Approach with Protein Force Field Description. J Chem Theory Comput 15, 5175–5193 (2019).

93. Lemkul, J. From Proteins to Perturbed Hamiltonians: A Suite of Tutorials for the GROMACS-2018 Molecular Simulation Package [Article v1.0]. Living J Comput Mol Sci 1, (2019).

94. Liebschner, D. et al. Macromolecular structure determination using X-rays, neutrons and electrons: recent developments in Phenix. Acta Crystallogr Sect D Struct Biology 75, 861–877 (2019).

95. Emsley, P., Lohkamp, B., Scott, W. G. & Cowtan, K. Features and development of Coot. Acta Crystallogr Sect D Biological Crystallogr 66, 486–501 (2010).

96. Williams, C. J. et al. MolProbity: More and better reference data for improved all-atom structure validation. Protein Sci 27, 293–315 (2018).

97. Pettersen, E. F. et al. UCSF ChimeraX: Structure visualization for researchers, educators, and developers. Protein Sci 30, 70–82 (2021).

98. Kidmose, R. T. et al. Namdinator – automatic molecular dynamics flexible fitting of structural models into cryo-EM and crystallography experimental maps. IUCrJ 6, 526–531 (2019).

