## Supplemental Figures for "Computational design of soluble functional analogues of integral membrane proteins"

### Supplementary information

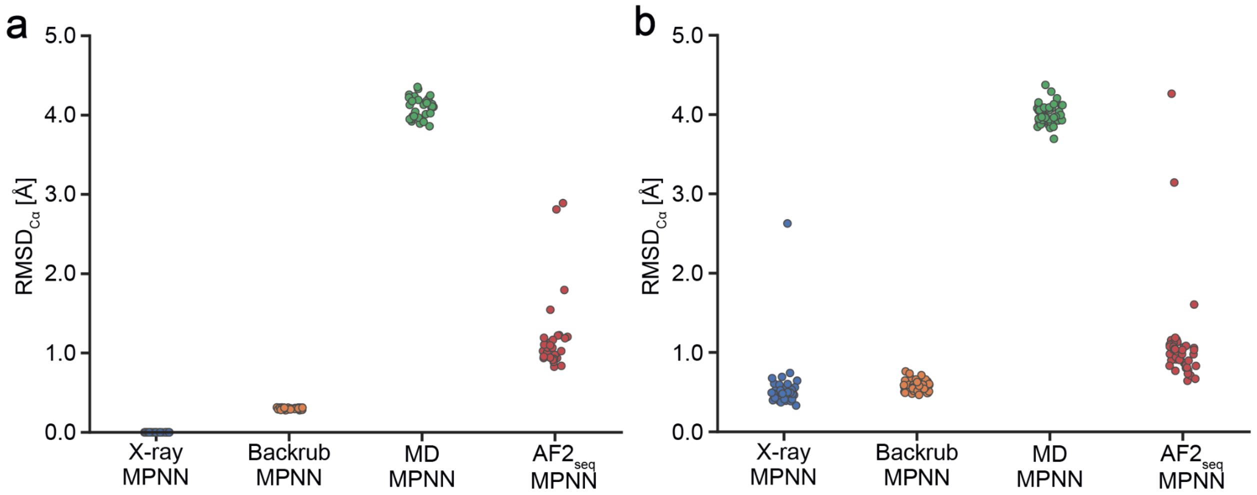


**Extended Data Fig. 1 | RMSD$_{C\alpha}$ of designed TIM-barrel structures vs the target fold crystal structure (PDB ID: 5BVL). a,** Backbone RMSD$_{C\alpha}$ deviations of input structures used for ProteinMPNN sequence redesign. **b,** Backbone RMSDCαdeviations of the highest ranked AF2 predicted structure derived from the ProteinMPNN-designed sequences from panel a.

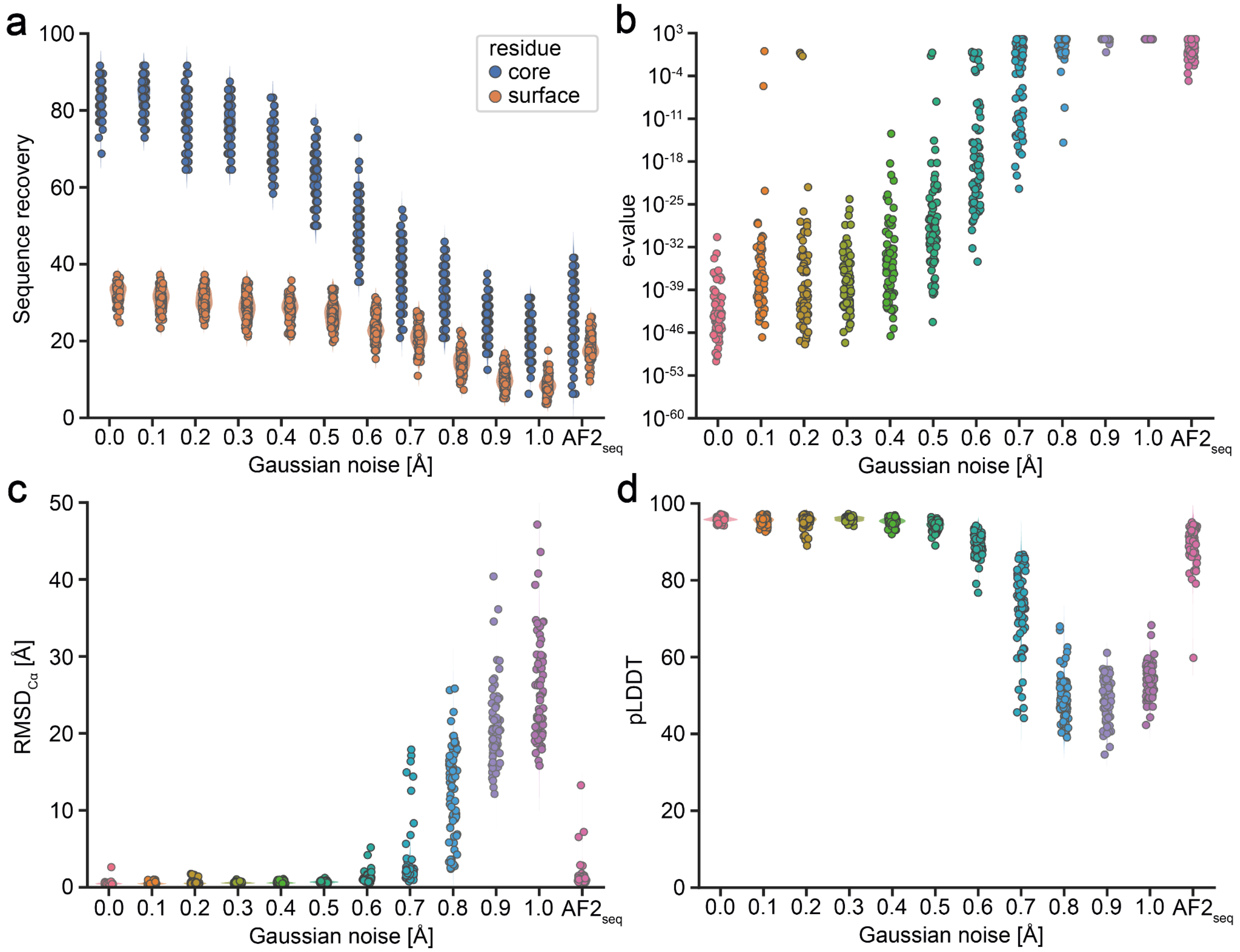


**Extended Data Fig. 2 | Computational analysis of the TIM-barrel fold (PDB ID: 5BVL) designs with Gaussian noise applied to the backbone coordinates.** ProteinMPNN designs with different values of Gaussian noise applied to the backbone atoms can generate feasible designs. They are not as diverse as AF2seq generated sequences as reflected by **a,** the percentage of sequence recovered in the core and on the surface and **b,** e-values of the generated sequences. The sequence diversity becomes comparable around 0.8 Å Gaussian noise, however, these designs have significantly higher **c,** backbone RMSD$_{C\alpha}$ deviations of output structures when the designed sequences are predicted using AF2 and lower **d,** AF2 confidence (pLDDT) scores.

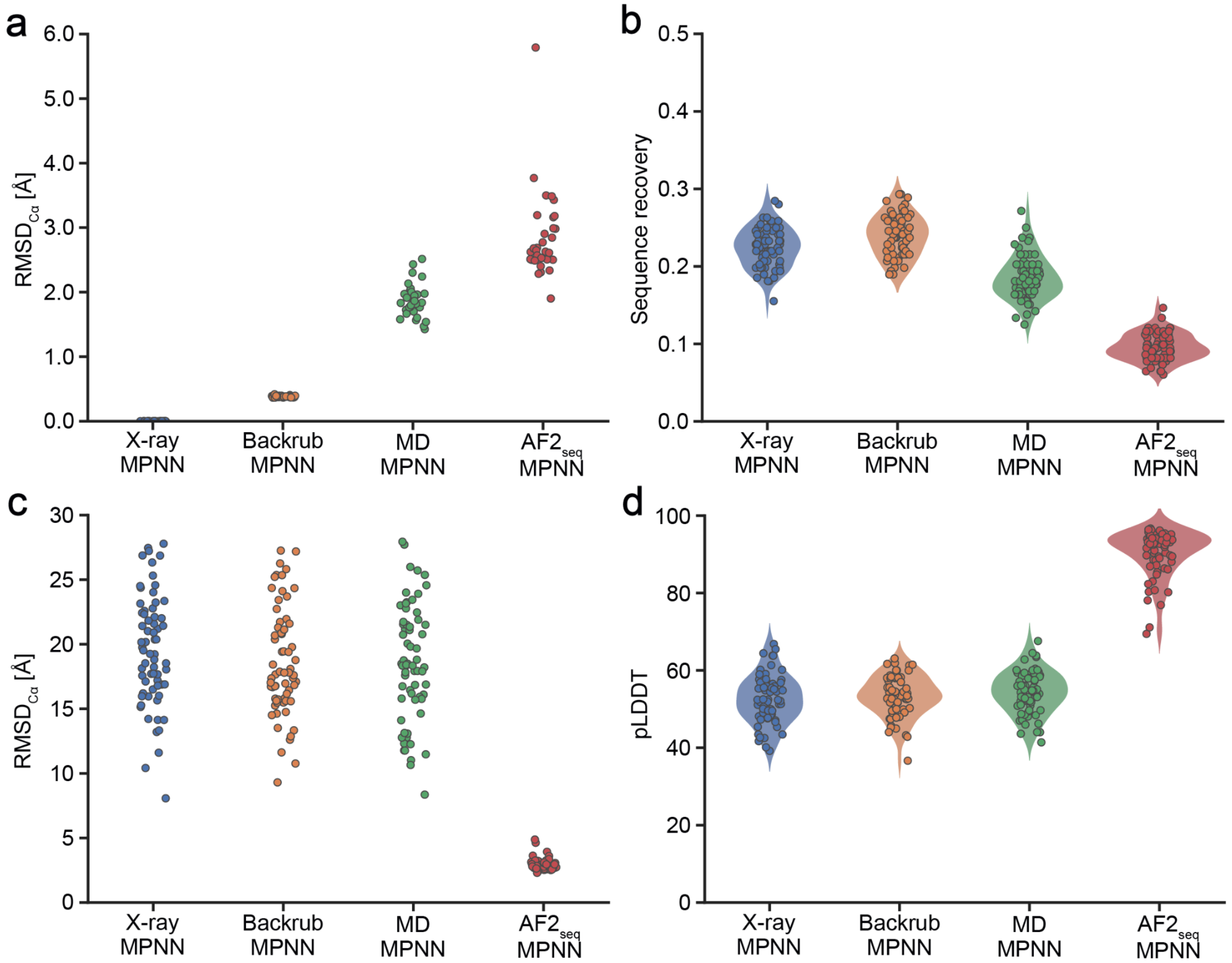


**Extended Data Fig. 3 | Computational analysis of GPCR backbone (PDB ID: 6FFI) perturbation methods for diversifying sequence design. a,** Backbone $RMSD_{C\alpha}$s relative to the reference crystal structure after being perturbed using Rosetta backrub, MD simulations, or using our $AF2_{seq}$ pipeline. **b,** Sequence recovery rates of ProteinMPNN sequence generation on the perturbed backbones. **c,** Backbone $RMSD_{C\alpha}$ of the top ranking AF2 model of ProteinMPNN sequences relative to the reference crystal structure. **d,** AF2 confidence scores of top ranking ProteinMPNN-derived predictions.

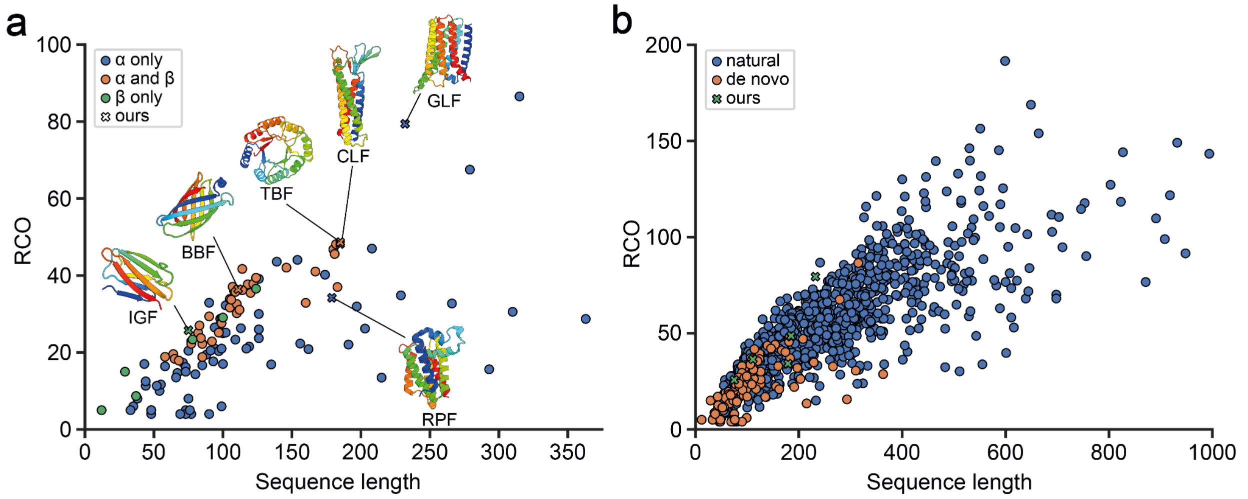


**Extended Data Fig. 4 | Relative Contact Order (RCO) plotted against sequence length of de novo designed and natural proteins.** Both RCO and sequence length describe the complexity of a protein (see Methods). This metric quantifies the number of contacts in a protein structure dependent on the sequence separation in order to capture the nonlocality in sequence of those contacts. **a,** Curated set of structures from computationally designed proteins reported by Verkuil et al.[41] and Woolfson[68] (shown in circles) were compared to the design targets in this paper (shown in crosses). This assessment shows that many of the designed topologies show high contact orders relative to other computationally designed proteins previously reported. Symbols are colored according to **b,** Comparison with natural folds shows that in general native proteins have higher contact orders then computationally designed proteins.

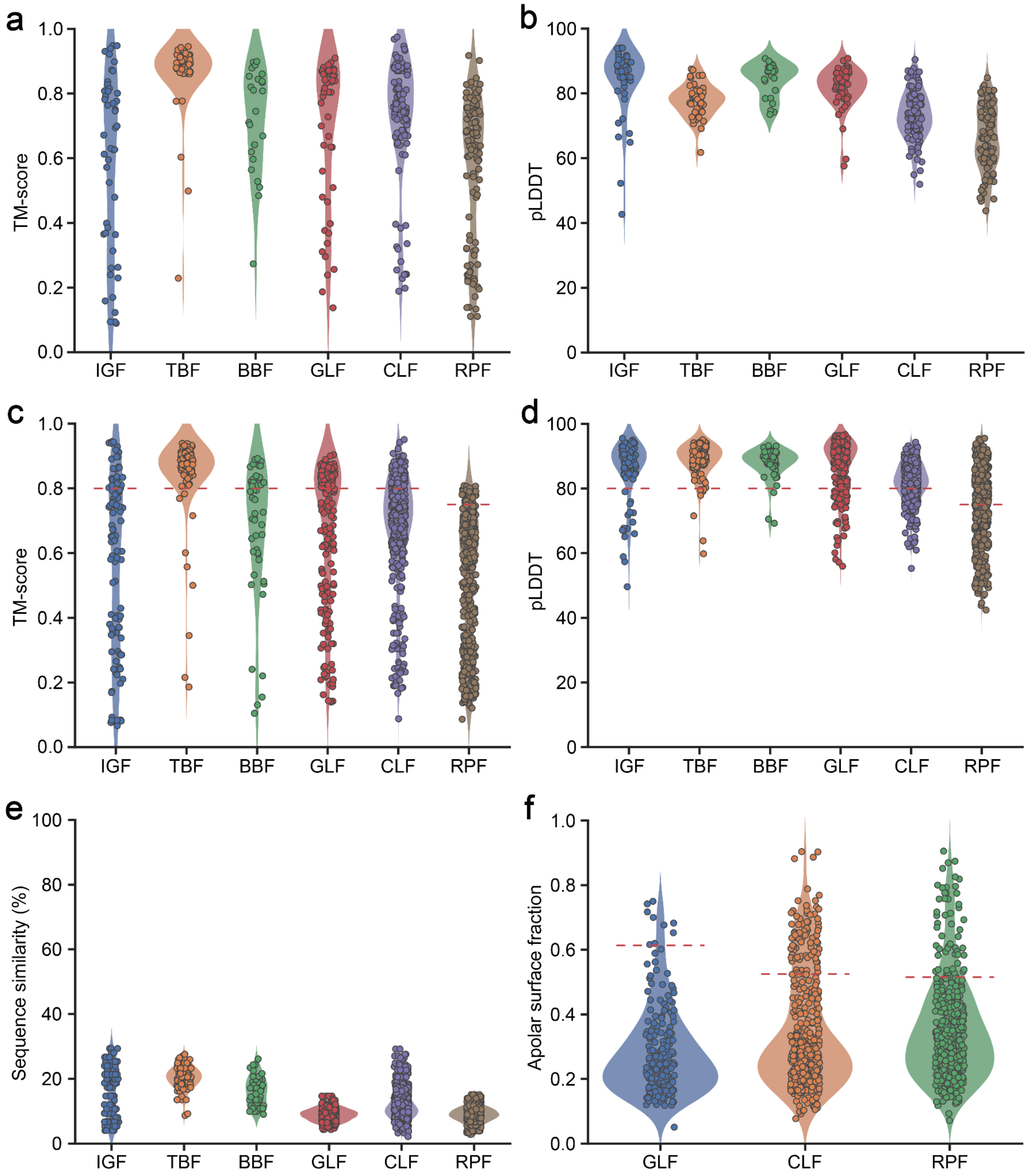


**Extended Data Fig. 5 | In silico analysis of the design folds.** **a**, shows the TM-scores of and **b**, confidence of the $AF2_{seq}$ generated sequences. The Ig-like fold (IGF), TIM-barrel fold (TBF) and β-barrel fold (BBF) are designed with proteinMPNN whilst the GPCR-like fold (GLF), claudin like fold (CLF) and rhomboid protease fold (RPF) are designed with proteinMPNN version trained on soluble proteins ($MPNN_{sol}$). The models of the $AF2_{seq}$-MPNN sequences are predicted using AF2 and the **c**, TM-scores relative to the designed model and **d**, confidence scores are shown. The dotted line depicts the cutoff values used for in vitro validation filtering. **e**, Plots the sequence similarity between the $AF2_{seq}$-MPNN designed sequence and original design target sequence. **f**, Fraction of apolar surface residues of the $AF2_{seq}$-$MPNN_{sol}$ designs. The dotted line represents the apolar surface fraction of each of the membrane protein design targets.

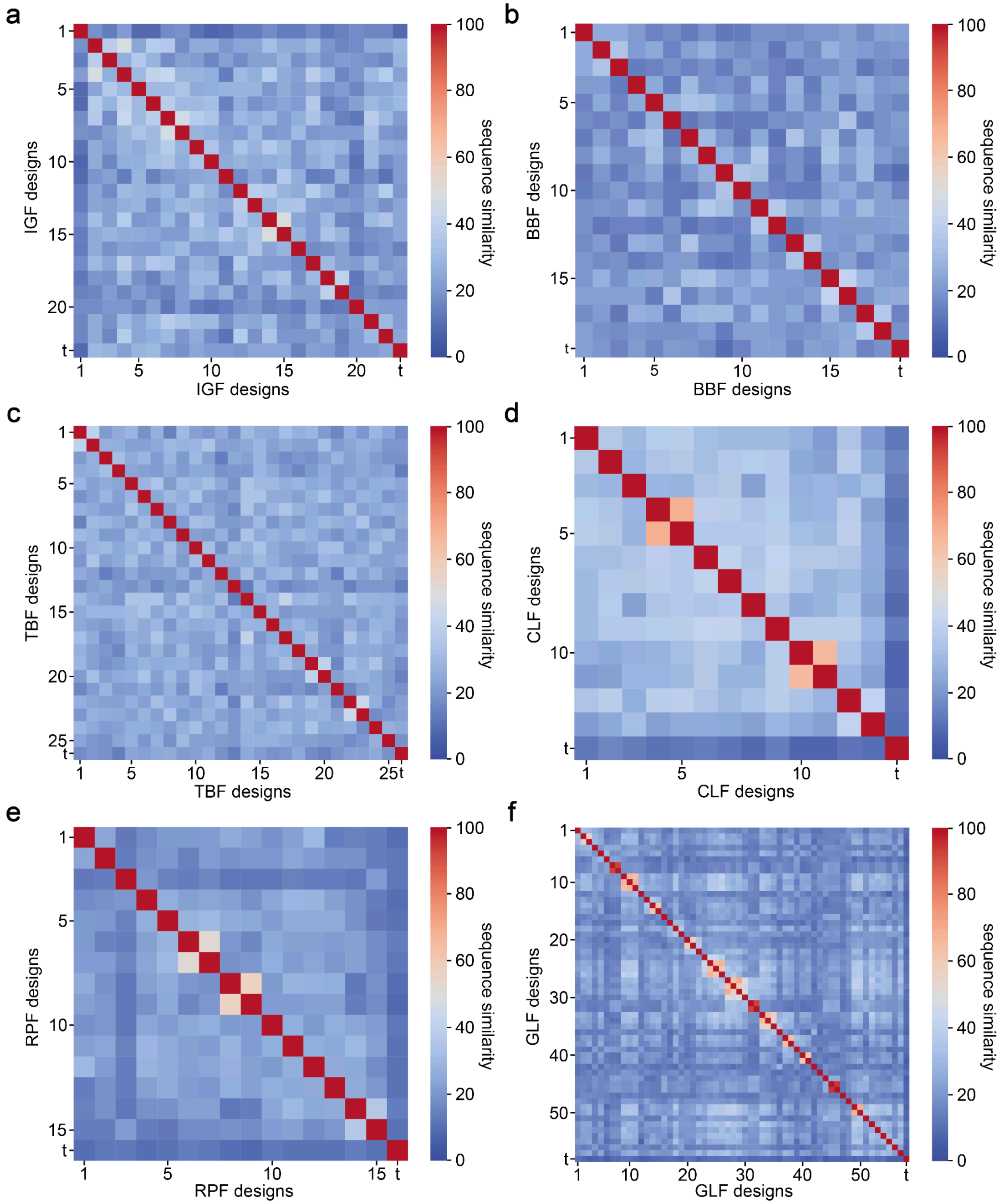


**Extended Data Fig. 6 | Sequence diversity of the generated designs.** Pairwise sequence similarities between **a,** Ig-Like Folds (IGF), **b,** β-barrel Folds (BBF), **c,** TIM-barrel Folds (TBF), **d,** Claudin-Like Folds (CLF), **e,** Rhomboid Protease Folds (RPF), and **f,** GPCR-Like Folds (GLF).

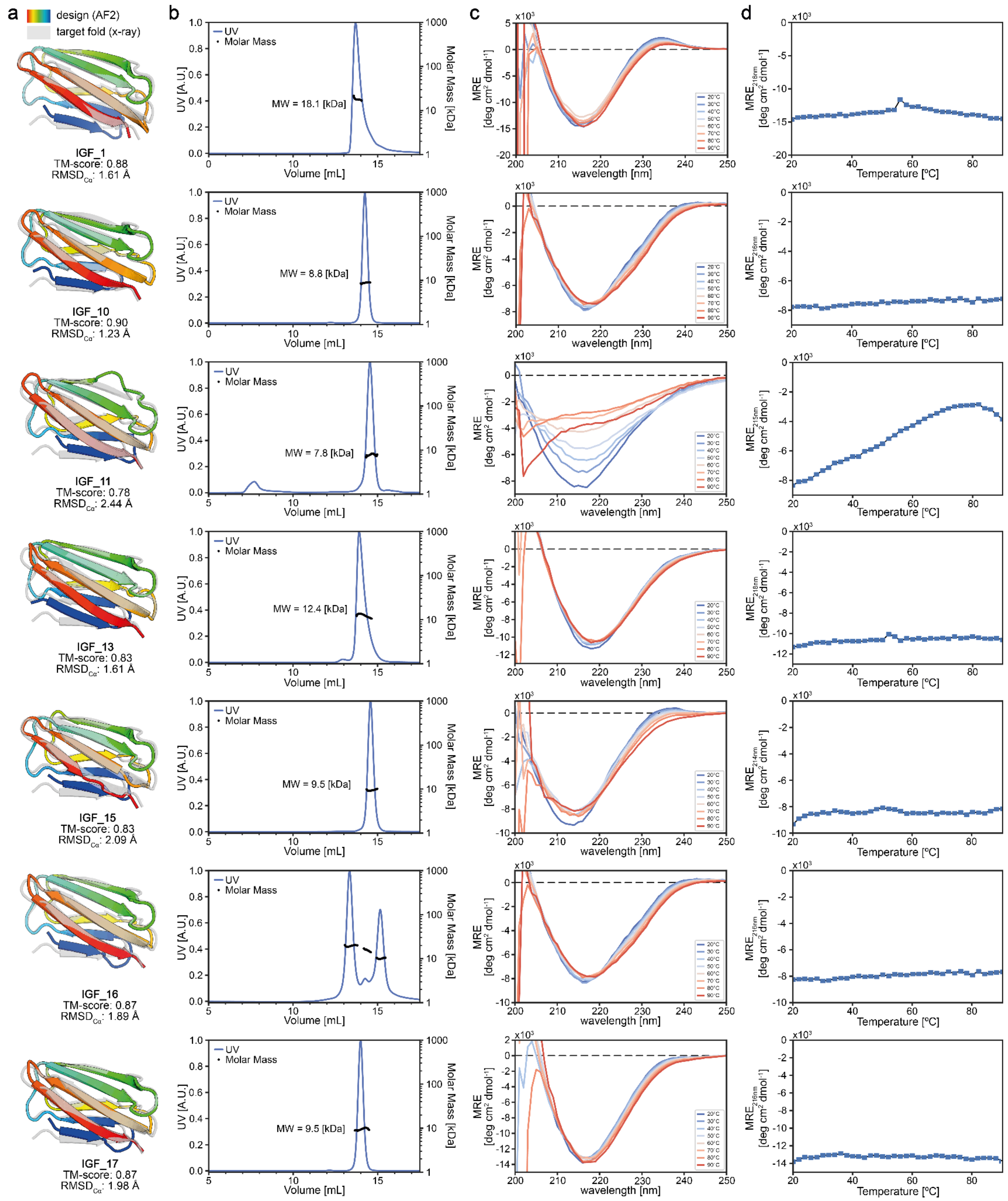


**Extended Data Fig. 7 | Biophysical characterization of designed Ig-like folds (IGF). a,** Cartoon depiction of design (colored) overlaid on the target fold (gray). **b,** SEC-MALS analysis of corresponding design in panel **a**. The expected Mw for the monomeric design ranges from 8.3 to 9.6 kDa. **c,** CD spectroscopy measurements at different temperatures. **d,** Thermostability based on CD measurement.

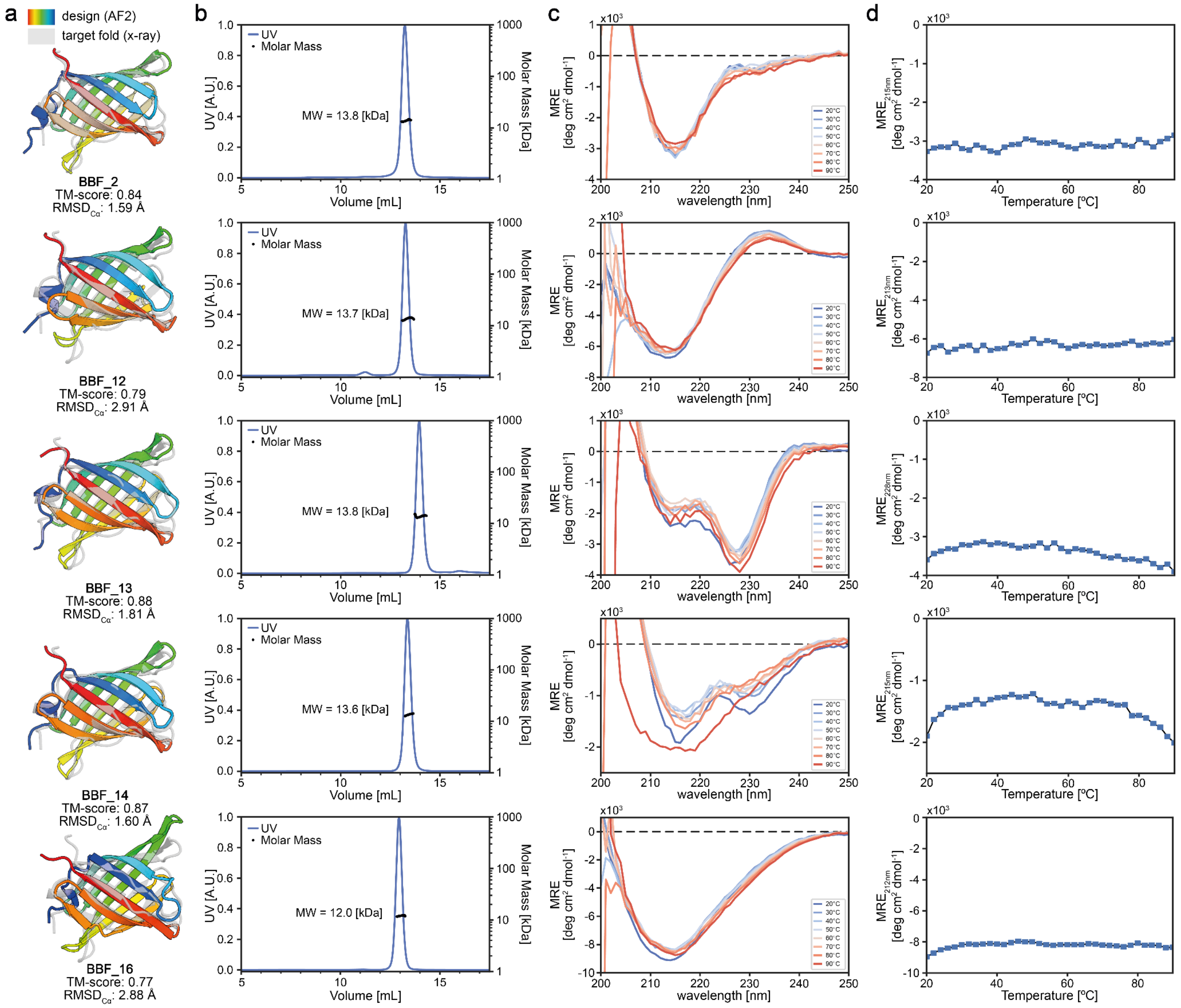


**Extended Data Fig. 8 | Biophysical characterization of designed β-barrel folds (BBF). a,** Cartoon depiction of design (colored) overlaid on the target fold (gray). **b,** SEC-MALS analysis of corresponding design in panel a. The expected Mw for the monomeric design ranges from 12.6 to 13.3 kDa. **c,** CD spectroscopy measurements at different temperatures. BBF_13 and BBF_14 present significant differences in their CD spectra as compared to the expected spectrum of the folded target structure. **d,** Thermostability curve based on CD measurement.

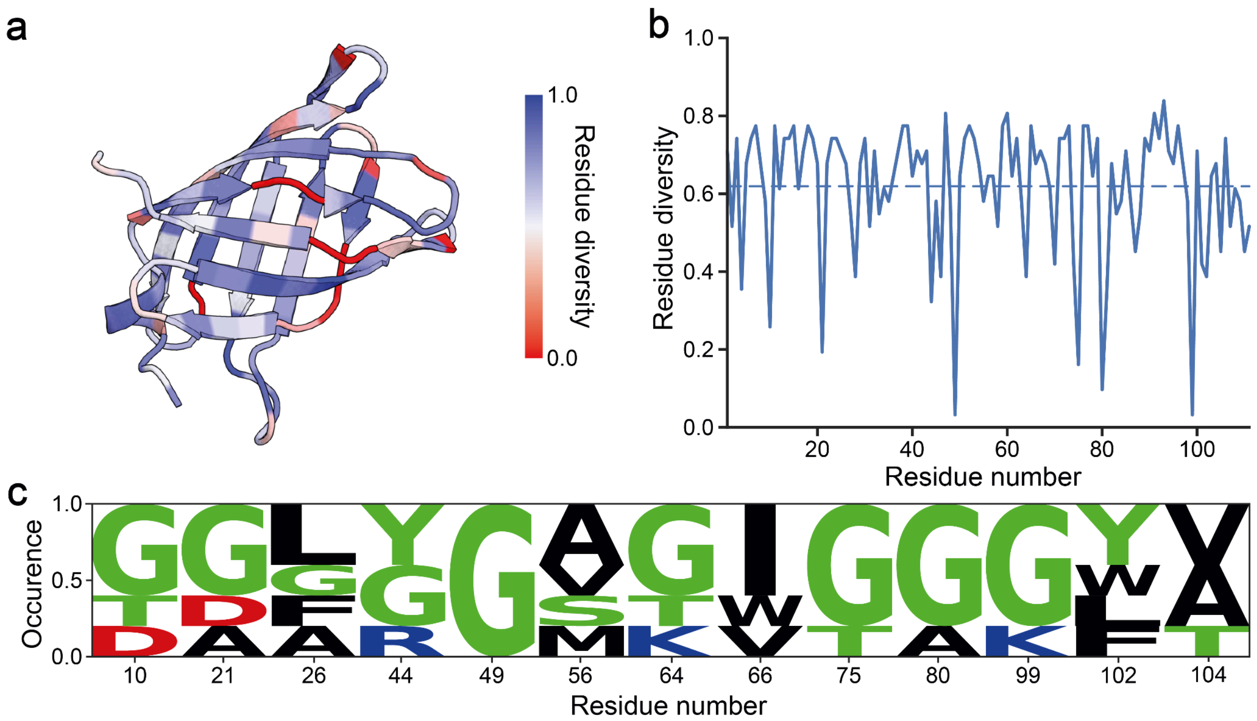


**Extended Data Fig. 9 | Sequence conservation analysis of β-barrel folds (BBF). a,** Cartoon depiction of example BBF design colored by sequence diversity of all designs on an individual residue level. **b,** Sequence diversity plotted on a per-residue level of all BBF designs. The dotted line represents the mean sequence diversity of the structure. **c,** Logoplot of residue occurrence of experimentally validated and folded BBF designs highlighting residue variability at sites critical for maintaining β-barrel topology.

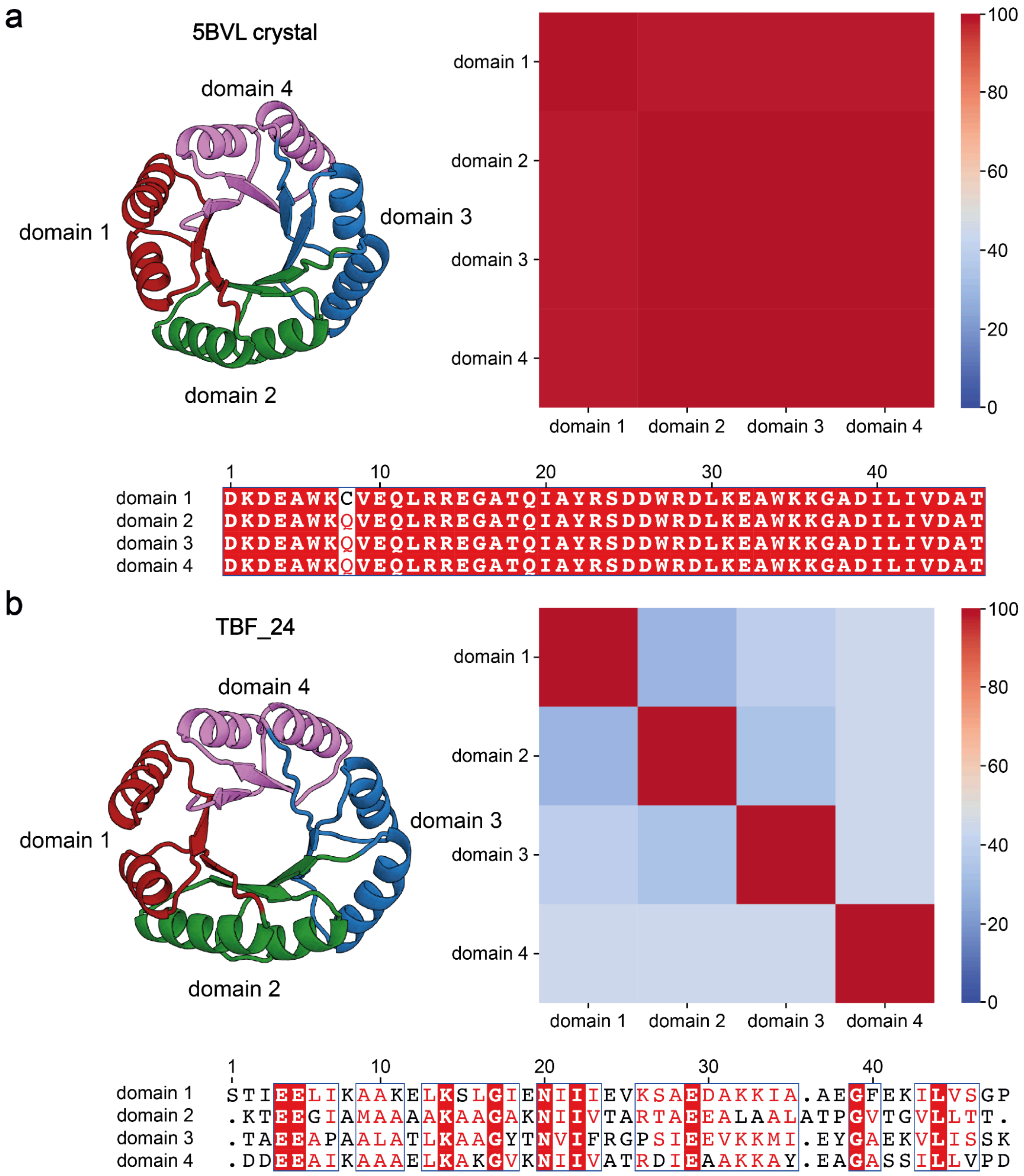


**Extended Data Fig. 10 | Sequence comparison of *de novo* designed TIM barrels. a,** Sequence similarity between the TIM barrel domain segments of the 4-fold symmetric design by Huang et al. **b,** Sequence similarity of the design TBF_24 without symmetric constraints. **a, b,** Crystal structure and sub-domain mapping on the left and sequence identity of the domains on the right.

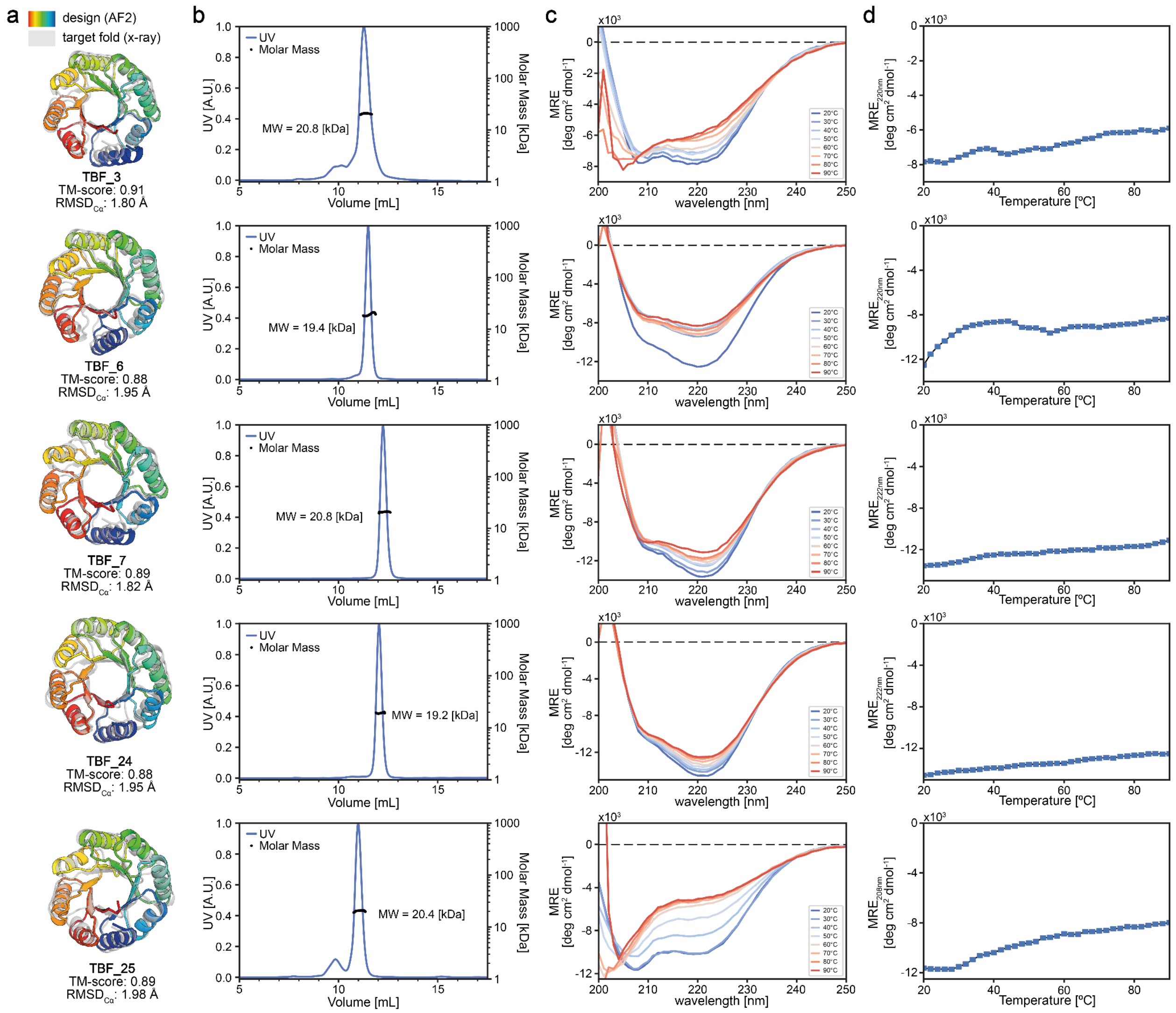


**Extended Data Fig. 11 | Biophysical characterization of designed TIM-barrel folds (TBF). a,** Cartoon depiction of design (colored) overlaid on the target fold (gray). **b,** SEC-MALS analysis of corresponding design in panel a. The expected Mw for the monomeric design ranges from 20.6 to 21.7 kDa. **c,** CD spectroscopy measurements at different temperatures. TBF_13 and TBF_25 present significant differences in their CD spectra as compared to the expected spectrum of the folded target structure. **d,** Thermostability curve based on CD measurement.

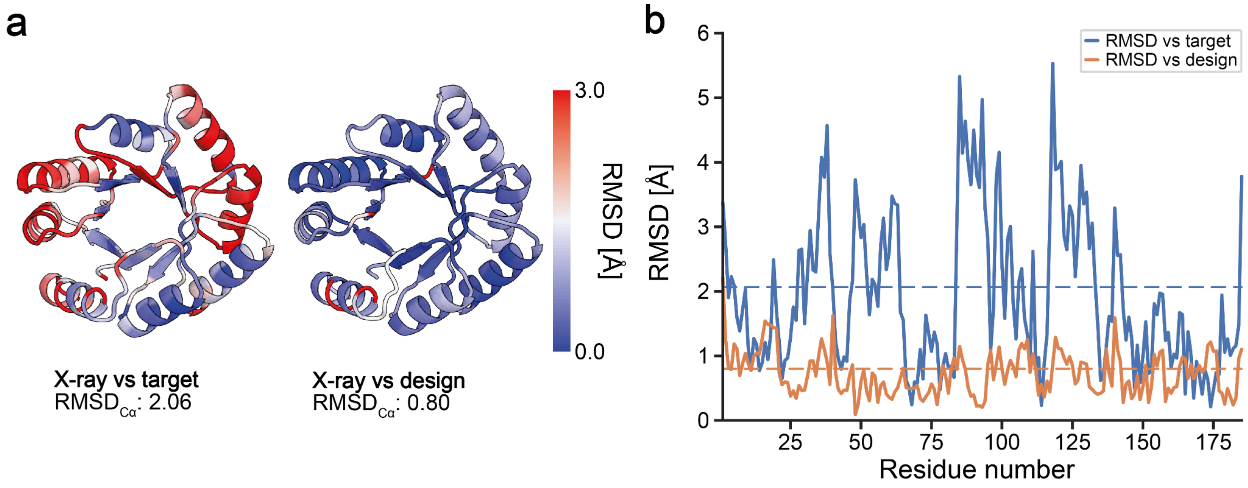


**Extended Data Fig. 12 | Backbone RMSD$_{Cα}$s of TBF_24 crystal structure relative to the target fold and design model. a,** Cartoon depiction of the crystal structure of TBF_24 colored by backbone RMSD$_{Cα}$ per residue when compared against the target fold (left) and the design model (right), with RMSD$_{Cα}$ per residue values plotted in panel **b.**

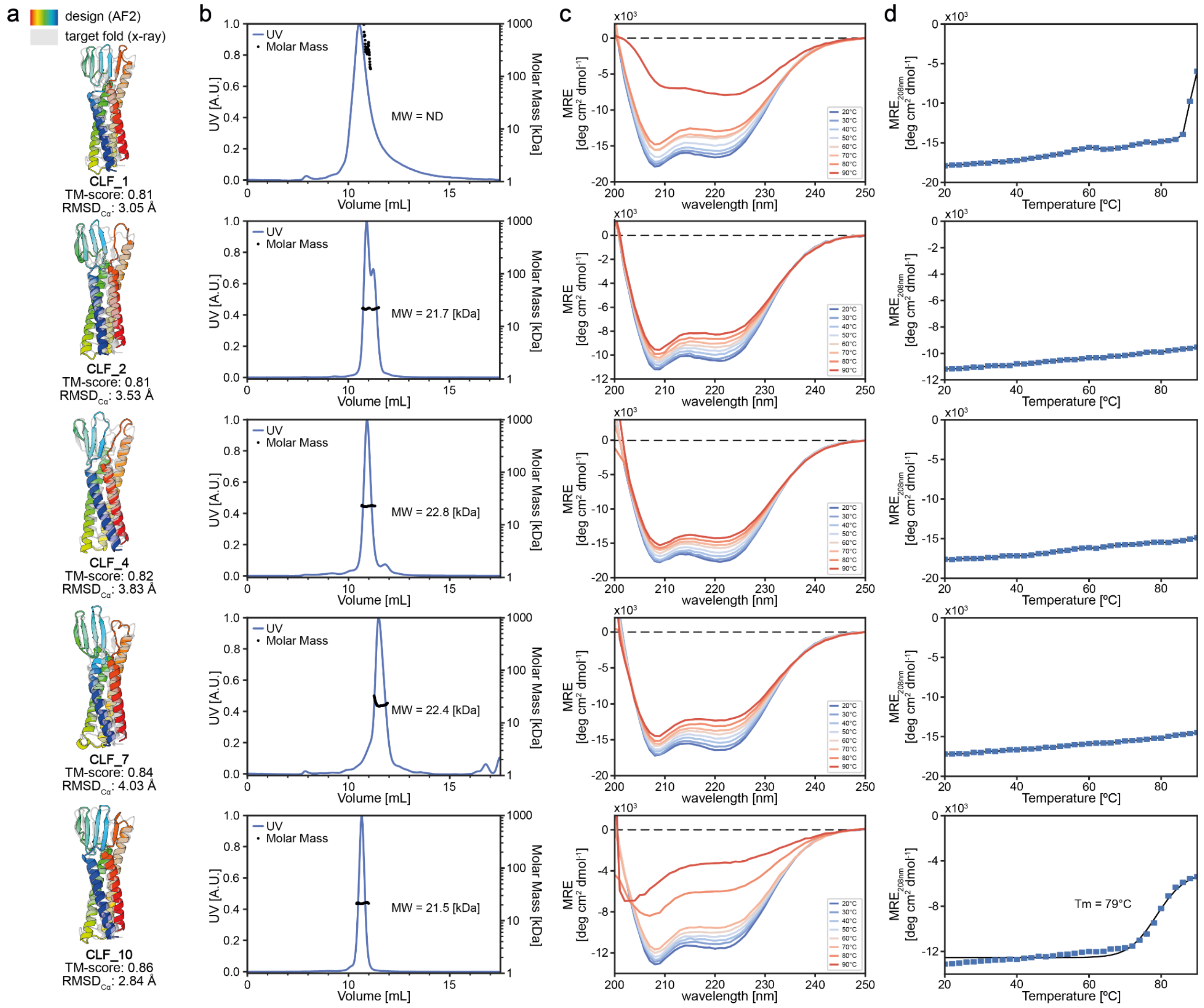


**Extended Data Fig. 13 | Biophysical characterization of designed Claudin-like folds (CLF). a,** Cartoon depiction of design (colored) overlaid on the target fold (gray). **b,** SEC-MALS analysis of corresponding design in panel a. The expected Mw for the monomeric design ranges from 21.5 to 22.5 kDa. **c,** CD spectroscopy measurements at different temperatures. **d,** Thermostability curve based on CD measurement.

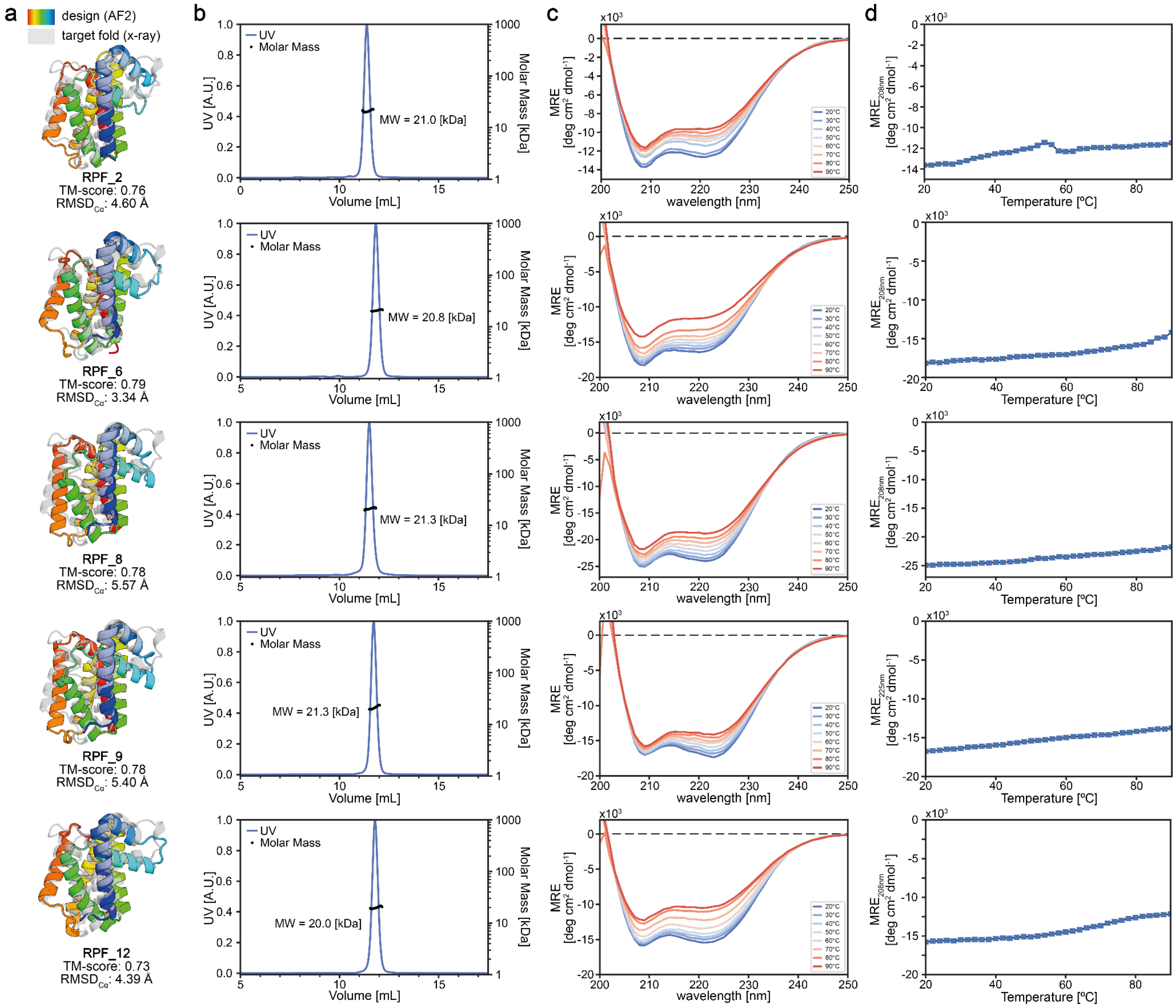


**Extended Data Fig. 14 | Biophysical characterization of designed Rhomboid protease folds (RPF).** **a,** Cartoon depiction of design (colored) overlaid on the target fold (gray). **b,** SEC-MALS analysis of corresponding design in panel a. The expected Mw for the monomeric design ranges from 20.7 to 21.8 kDa. **c,** CD spectroscopy measurements at different temperatures. **d,** Thermostability curve based on CD measurement.

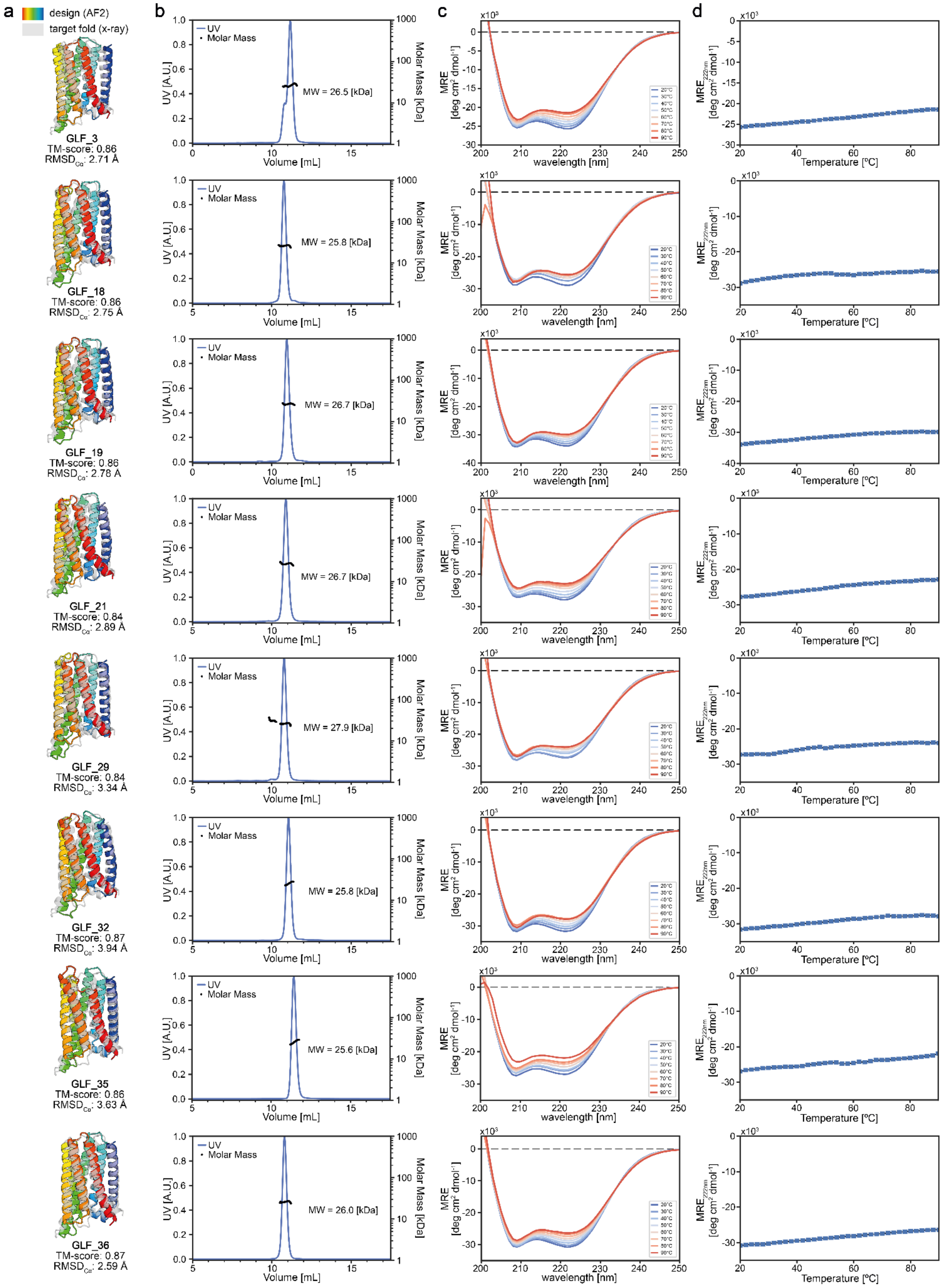
a
design (AF2)
target fold (x-ray)
b
c
d
GLF_3
TM-score: 0.86
RMSD: 2.71 Å
MW = 26.5 [kDa]
GLF_18
TM-score: 0.86
RMSD: 2.75 Å
MW = 25.8 [kDa]
GLF_19
TM-score: 0.86
RMSD: 2.78 Å
MW = 26.7 [kDa]
GLF_21
TM-score: 0.84
RMSD: 2.89 Å
MW = 26.7 [kDa]
GLF_29
TM-score: 0.84
RMSD: 3.34 Å
MW = 27.9 [kDa]
GLF_32
TM-score: 0.87
RMSD: 3.94 Å
MW = 25.8 [kDa]
GLF_35
TM-score: 0.86
RMSD: 3.63 Å
MW = 25.6 [kDa]
GLF_36
TM-score: 0.87
RMSD: 2.59 Å
MW = 26.0 [kDa]
UV
Molar Mass
UV [A.U.]
Molar Mass [kDa]
Volume [mL]
MRE [deg cm² dmol⁻¹]
wavelength [nm]
Temperature [ºC]

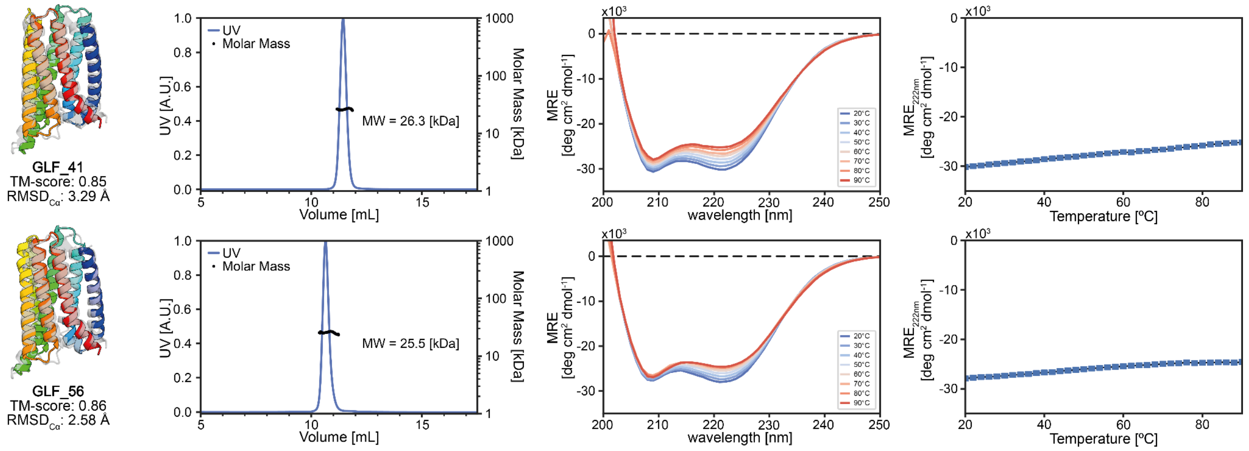


**Extended Data Fig. 15 | Biophysical characterization of designed GPCR-like folds (GLF). a,** Cartoon depiction of design (colored) overlaid on the target fold (gray). **b,** SEC-MALS analysis of corresponding design in panel a. The expected Mw for the monomeric design ranges from 27.2 to 28.1 kDa. **c,** CD spectroscopy measurements at different temperatures. **d,** Thermostability curve based on CD measurements.

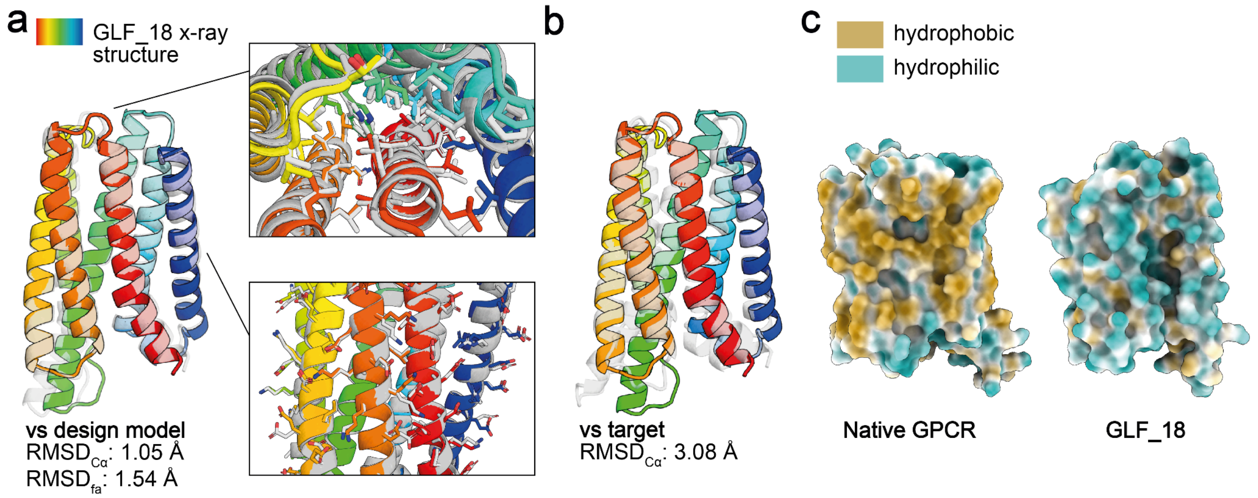


**Extended Data Fig. 16 | GLF_18 X-ray structure. a,** X-ray structure of GLF_18 (colored) superimposed on the design model (gray). **b,** X-ray structure of GLF_18 (colored) superimposed on the design model (gray). **c,** Molecular lipophilicity potential of the surface of the GPCR design target and the soluble GLF_18 design. After redesign of the original membrane folds with $MPNN_{sol}$, the hydrophobicity (yellow) of the surface is significantly reduced, and polarity increased (blue). $RMSD_{C\alpha}$ - root mean square deviation computed over the Cα atoms of the backbone. $RMSD_{fa}$ - root mean square deviation computed over all the atoms in the structure.

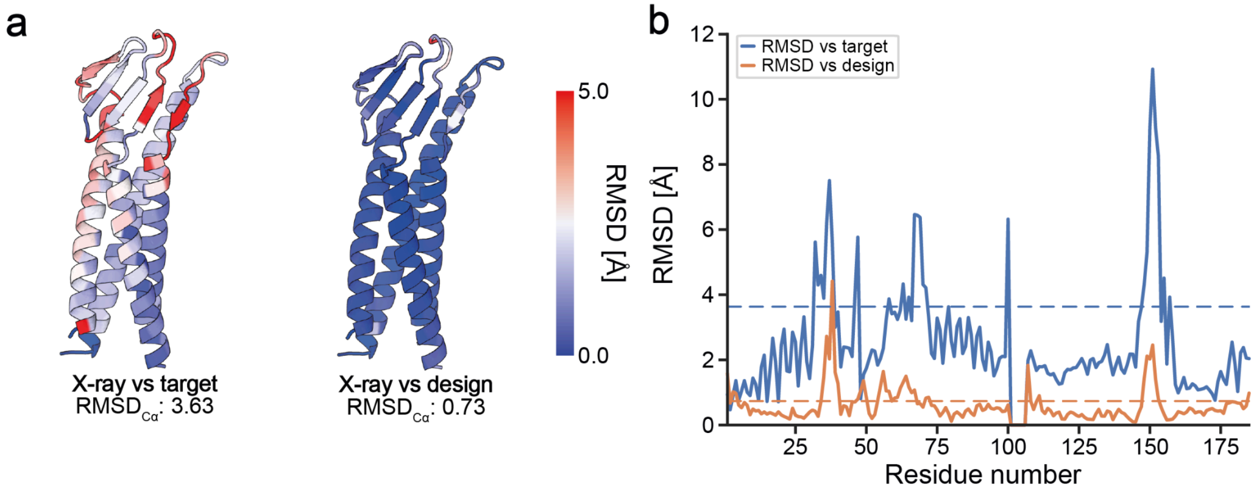


**Extended Data Fig. 17 | Backbone RMSD$_{C\alpha}$s of CLF_4 crystal structure relative to the target fold and design model.** **a**, cartoon representation of the crystal structure of CLF_4, with the left and right structures colored by backbone RMSD$_{C\alpha}$ per residue when compared against the target fold and the design model, respectively. Panel **b** displays the RMSD$_{C\alpha}$ per residue values for both comparisons. In the comparison with the target fold the largest differences are found in the β-sheet region.

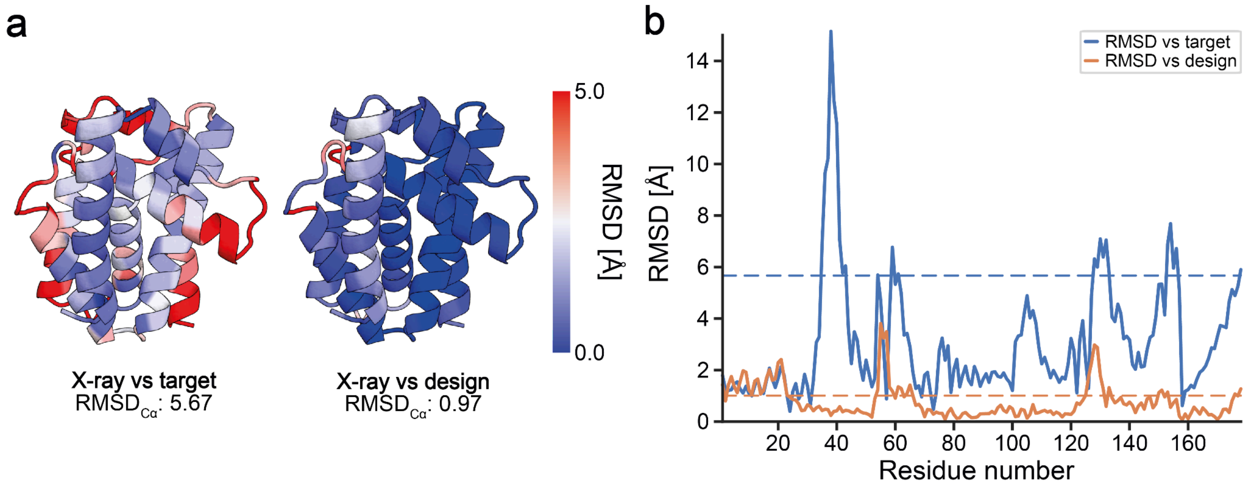


**Extended Data Fig. 18 | Backbone RMSD$_{C\alpha}$s of RPF_9 crystal structure relative to the target fold and design model.** **a**, cartoon representation of the crystal structure of RPF_9, with the left and right structures colored by backbone RMSD$_{C\alpha}$ per residue when compared against the target fold and the design model, respectively. Panel **b** displays the RMSD$_{C\alpha}$ per residue values for both comparisons. The loop region between residues 35 and 42 was not structurally similar between the RPF_9 design and the target fold, however, the X-ray structure closely matches the designed model.

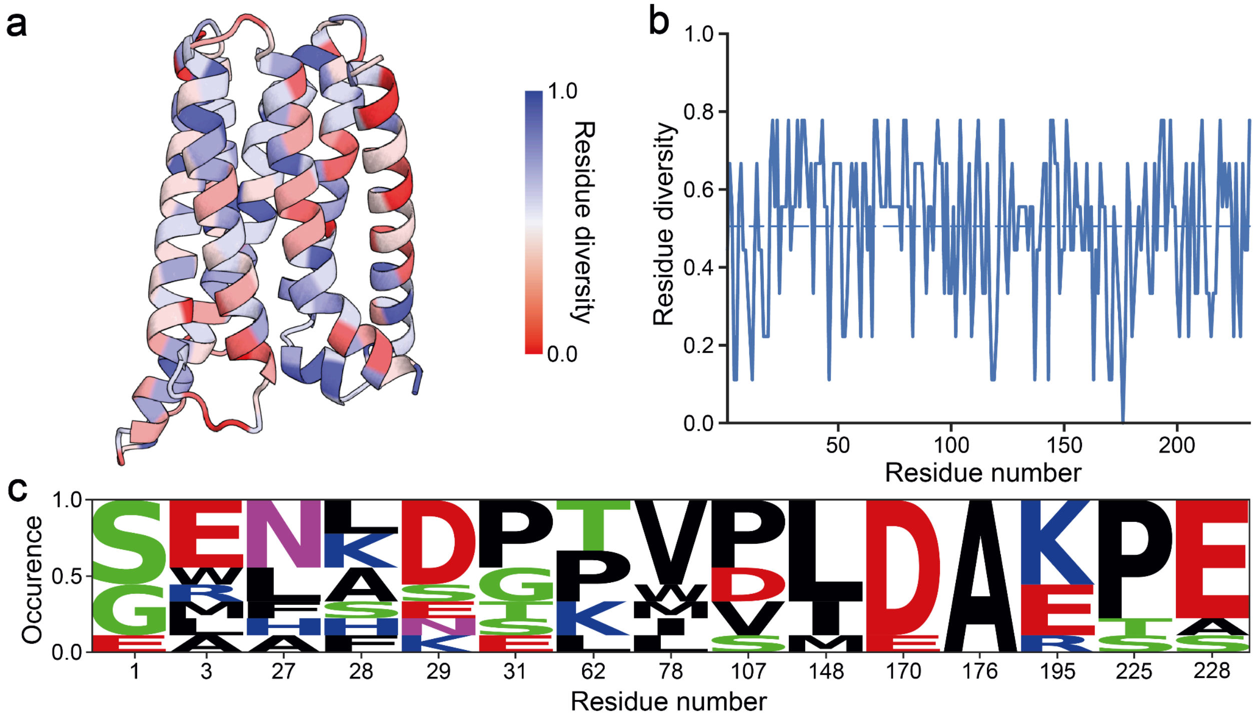


**Extended Data Fig. 19 | Sequence conservation analysis of GPCR-like folds (GLF). a**, Cartoon representation of a GLF colored by residue diversity of the in vitro validated folded designs. **b**, The sequence diversity of all GLF designs is visualized on a per-residue level in the plot, with a dotted line indicating the average sequence diversity across the structure. **c,** depiction of the occurrence of residues in experimentally validated and folded GLF designs. The natural GLF contained a highly conserved DRY motif in the first intracellular loop (residues 26 to 28) and a PXXY motif (residues 225 to 227) in the seventh helix which are not present in our designs. All other positions depicted contained proline residues in the design target, which have a higher prevalence for some positions but are not required.

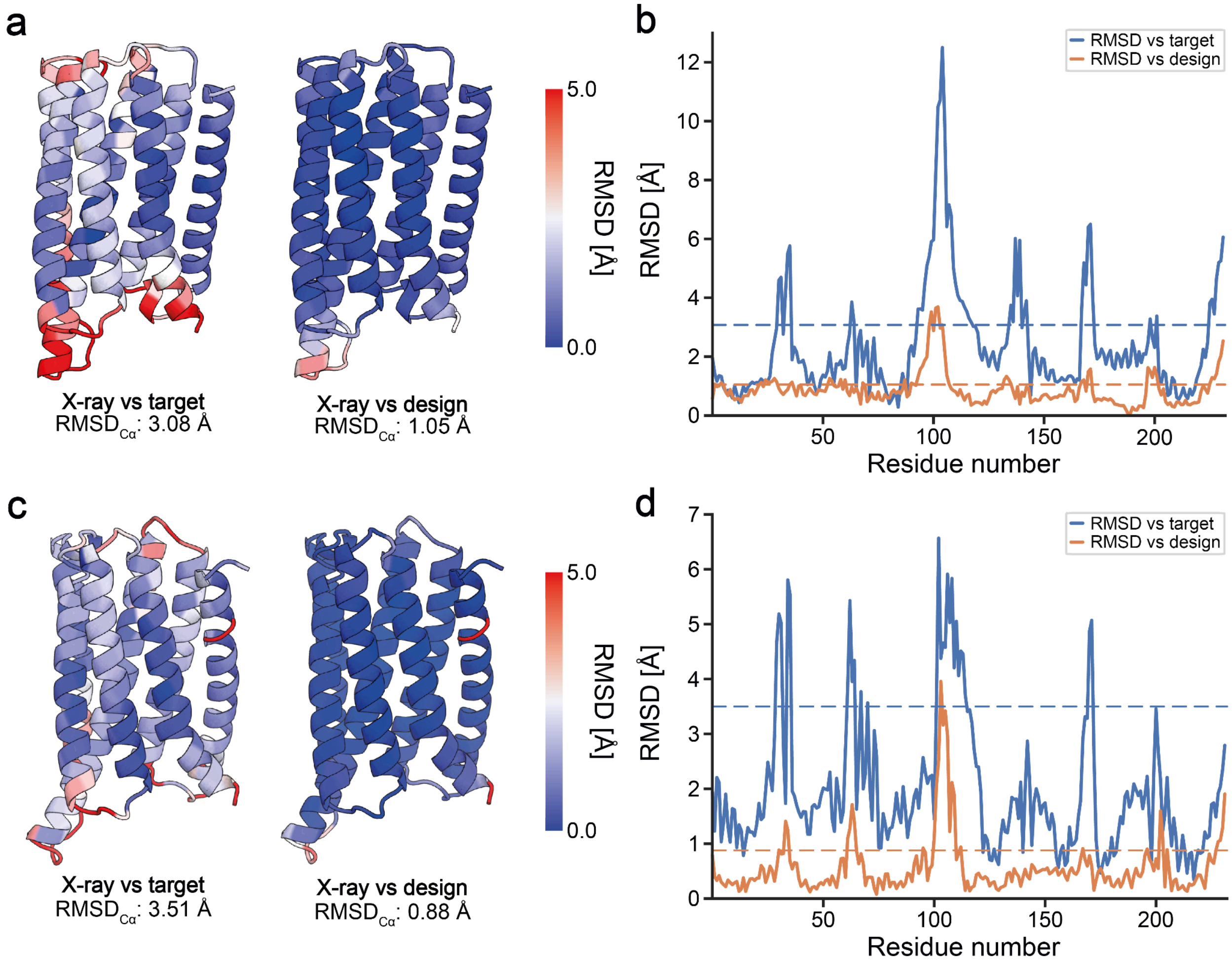


**Extended Data Fig. 20 | Backbone $RMSD_{C\alpha}$s of crystallized GLF_18 and GLF_32 crystal structures relative to the target fold and design models. a,** Cartoon depiction of the crystal structure of GLF_18 colored by backbone $RMSD_{C\alpha}$ per residue when compared against the target fold (left) and the design model (right), with individual values plotted in panel **b. c,** Depiction of the GLF_32 crystal structure colored by backbone $RMSD_{C\alpha}$, with individual values plotted per residue in panel **d.**

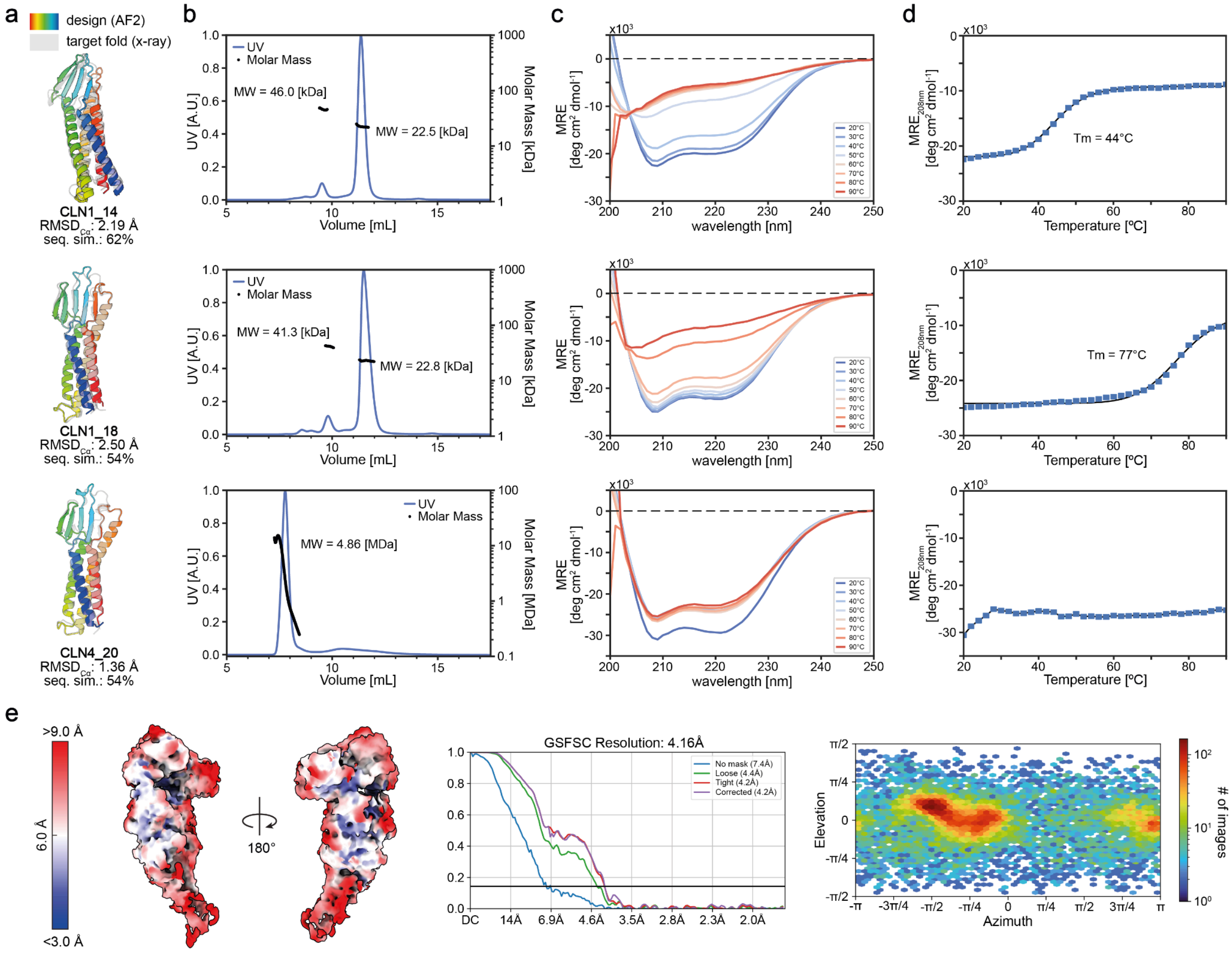


**Extended Data Fig. 21 | Biophysical characterization of the solubilized Human Claudin-1 (CLN1) and Claudin-4 (CLN4) presenting functional motifs. a,** Cartoon depiction of design (colored) overlaid on the target fold (gray). **b,** SEC-MALS analysis of corresponding design in panel a. The expected Mw for the monomeric design ranges from 27.2 to 28.1 kDa. **c,** CD spectroscopy measurements at different temperatures. **d,** Thermostability curve based on CD measurements. **e,** Left, CLN4_20-cCpE-COP2-nanobody complex unsharpened cryoEM map used for model docking colored by local resolution. Middle, Gold-standard FSC curve with resolution cutoff indicated at 0.143. Right, Particle distribution heatmap of the final reconstruction.

a

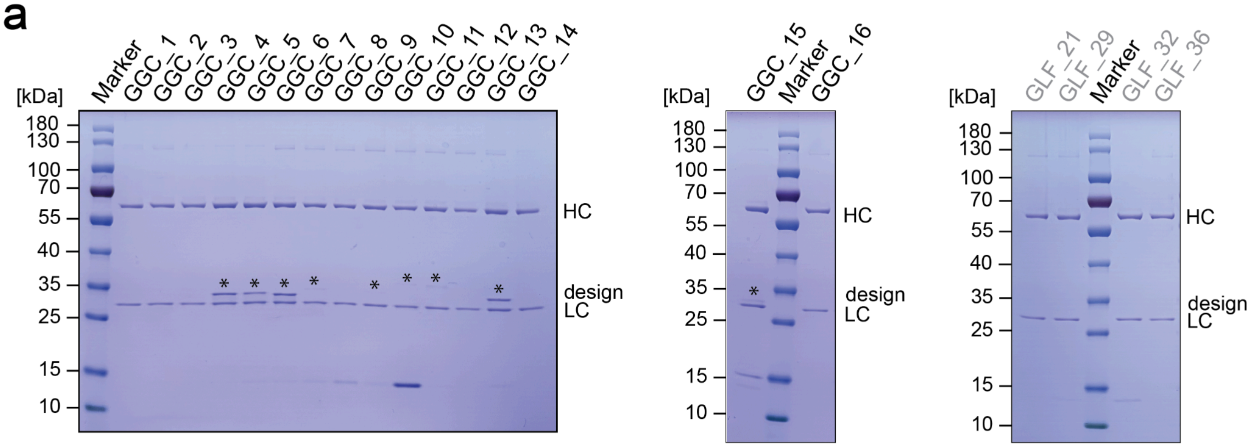


**Extended Data Fig. 22 | Pulldown screening of antibody binding to GGC designs. a,** SDS-PAGE analysis of eluted fractions from antibody binding screens to soluble GPCR scaffolds with transplanted ICL3 loops from Ghrelin GPCR receptor. Antibody light-chain (LC) and heavy-chain (HC) are highlighted, design corresponds to soluble GPCR construct. Pulled down constructs are highlighted by asterisks. Soluble scaffolds without transplanted loops are highlighted in gray and serve as negative controls.

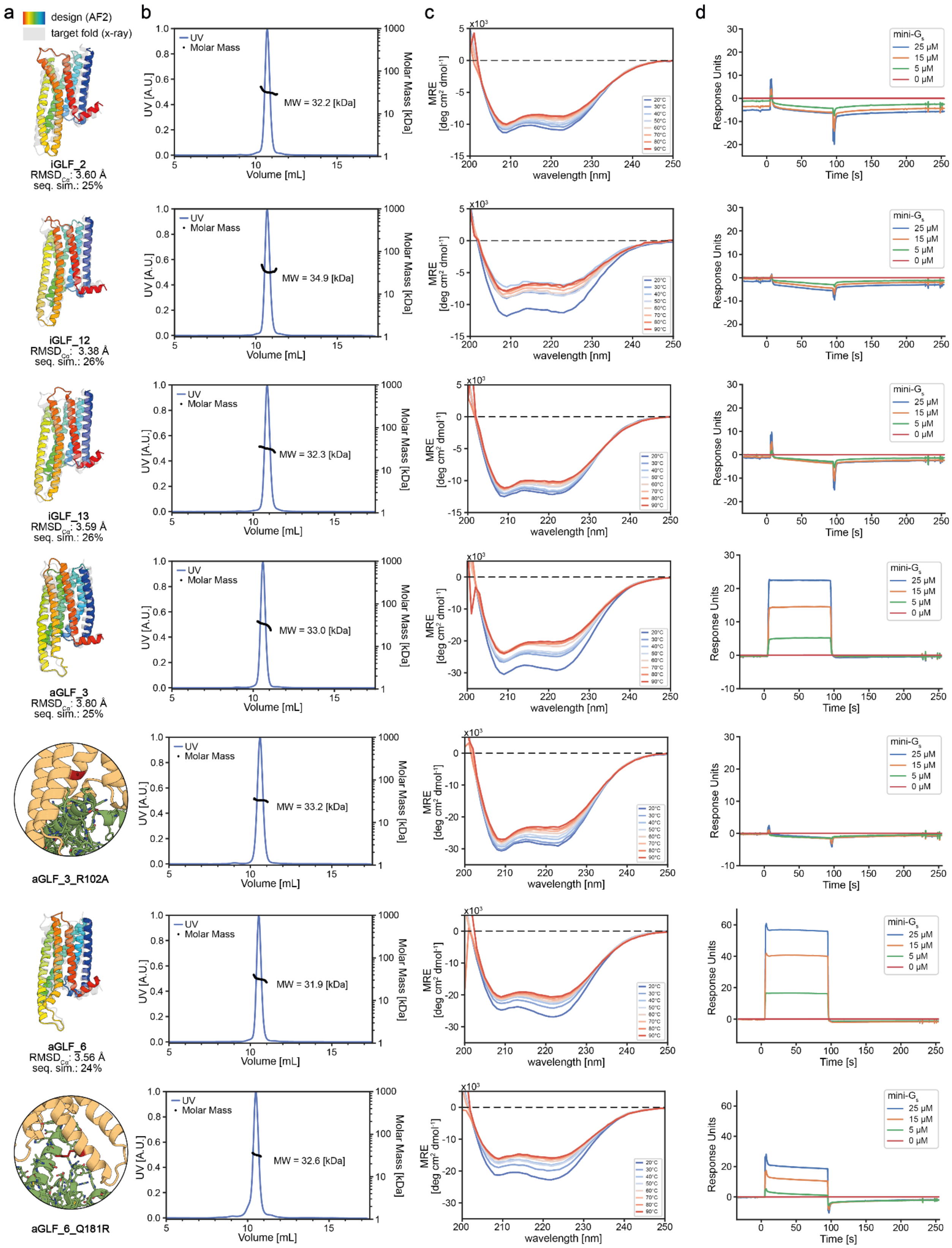
a
design (AF2)
target fold (x-ray)
b
c
d
iGLF_2
RMSD$_{C\alpha}$: 3.60 Å
seq. sim.: 25%
MW = 32.2 [kDa]
iGLF_12
RMSD$_{C\alpha}$: 3.38 Å
seq. sim.: 26%
MW = 34.9 [kDa]
iGLF_13
RMSD$_{C\alpha}$: 3.59 Å
seq. sim.: 26%
MW = 32.3 [kDa]
aGLF_3
RMSD$_{C\alpha}$: 3.80 Å
seq. sim.: 25%
MW = 33.0 [kDa]
aGLF_3_R102A
MW = 33.2 [kDa]
aGLF_6
RMSD$_{C\alpha}$: 3.56 Å
seq. sim.: 24%
MW = 31.9 [kDa]
aGLF_6_Q181R
MW = 32.6 [kDa]
UV
Molar Mass
UV [A.U.]
Molar Mass [kDa]
Volume [mL]
MRE [deg cm$^2$ dmol$^{-1}$]
x10$^3$
wavelength [nm]
20°C
30°C
40°C
50°C
60°C
70°C
80°C
90°C
mini-G$_s$
25 µM
15 µM
5 µM
0 µM
Response Units
Time [s]

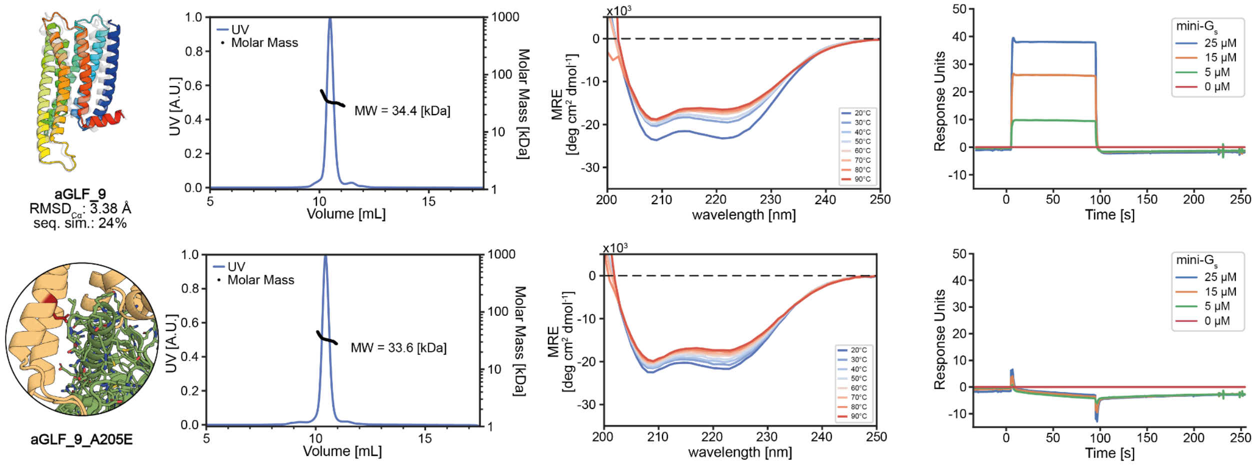


**Extended Data Fig. 23 | Biophysical characterization of the conformational specific GPCR-like fold analogs containing g-protein binding sites. a,** Cartoon depiction of design (colored) overlaid on the target fold (gray). The point mutants of the aGLF (orange) show a zoom with the point mutation (red) in the presence of mini-$g_s$ (green). **b,** SEC-MALS analysis of corresponding design in panel a. The expected Mw for the monomeric design ranges from 33.1 to 34.1 kDa. **c,** SPR sensorgram of different mini-$g_s$ concentrations in the presence of the designs from panel a.

**Supplementary Table S1| Crystallographic data collection and refinement statistics.**

| | *TBF_24 (PDB: 8OYS)* | *CLF_4 (PDB: 8OYV)* | *RPF_9 (PDB: 8OYW)* | *GLF_18 (PDB: 8OYX)* | *GLF_32 (PDB: 8OYY)* |
|---|---|---|---|---|---|
| **Data collection** | | | | | |
| Space group | $P\ 2_1\ 2_1\ 2_1$ | P 1 | $P\ 2_1\ 2_1\ 2_1$ | P 1 | $P\ 2_1\ 2_1\ 2_1$ |
| Cell dimensions | | | | | |
| *a*, *b*, *c* (Å) | 36.63, 44.41, 125.46 | 35.56, 50.77, 54.93 | 39.72, 66.68, 73.39 | 37.73, 52.08, 58.39 | 39.28, 56.78, 202.97 |
| a, b, g (°) | 90.00, 90.00, 90.00 | 107.61, 90.50, 90.06 | 90.00, 90.00, 90.00 | 94.08, 104.36, 110.80 | 90.00, 90.00, 90.00 |
| Wavelength (Å) | 1.000 | 1.000 | 1.000 | 1.000 | 0.966 |
| Resolution (Å) | 62.73 - 1.34 (1.39 - 1.34) | 48.39 - 2.80 (2.90 - 2.80) | 34.93 - 1.50 (1.55 - 1.50) | 39.97 - 2.11 (2.19 - 2.11) | 49.55 - 1.85 (1.92 - 1.85) |
| Unique reflect. | 46889 (4601) | 8007 (858) | 31885 (3164) | 20175 (2144) | 38572 (3930) |
| $R_{merge}$ | 0.058 (1.422) | 0.1069 (1.588) | 0.0805 (1.196) | 0.050 (0.335) | 0.096 (1.526) |
| *I* / s*I* | 15.6 (1.4) | 6.34 (1.42) | 11.8 (1.5) | 9.9 (3.4) | 15.4 (1.7) |
| Complete. (%) | 99.9 (100.0) | 88.0 (95.1) | 99.6 (99.7) | 88.5 (92.8) | 97.7 (99.8) |
| Redundancy | 9.2 (9.5) | 2.5 (2.7) | 8.1 (7.5) | 1.7 (1.7) | 11.6 (11.6) |
| **Refinement** | | | | | |
| Resolution (Å) | 62.73 - 1.34 | 48.39 - 2.80 | 34.93 - 1.5 | 39.97 - 2.11 | 49.55 - 1.85 |
| No. reflections | 46880 (4599) | 7983 (858) | 31846 (3158) | 20185 (2144) | 38882 (3924) |
| $R_{work}$ / $R_{free}$ | 0.1673/0.1885 | 0.2725/0.3168 | 0.1967/0.2159 | 0.2021/0.2500 | 0.1936/0.2379 |

| | | | | | |
|---|---|---|---|---|---|
| **No. atoms** | | | | | |
| Protein | 1390 | 2950 | 1502 | 3686 | 3749 |
| Ligand/ion | 2 | 0 | 1 | 10 | 6 |
| Water | 232 | 1 | 163 | 44 | 203 |
| ***B*-factors** | | | | | |
| Protein | 28.5 | 97.7 | 38.6 | 53.5 | 41.1 |
| Ligand/ion | 46.8 | - | 55.1 | 74.7 | 58.4 |
| Water | 39.1 | 74.0 | 43.2 | 49.2 | 42.4 |
| **R.m.s. deviations** | | | | | |
| Bond lengths (Å) | 0.006 | 0.002 | 0.008 | 0.006 | 0.006 |
| Bond angles (°) | 0.840 | 0.480 | 0.680 | 0.880 | 0.750 |
| **Ramachandran plot** | | | | | |
| Favored (%) | 93.99 | 97.45 | 98.86 | 98.67 | 100.00 |
| Allowed (%) | 6.01 | 2.55 | 1.14 | 1.33 | 0.00 |
| Outliers (%) | 0.00 | 0.00 | 0.00 | 0.00 | 0.00 |
| **Rotamer outliers (%)** | 0.00 | 0.91 | 1.90 | 0.00 | 1.02 |
| **Clashscore** | 1.39 | 3.51 | 1.34 | 6.12 | 3.01 |

*Values in parentheses are for the highest-resolution shell.

**Supplementary Table S2| Cryo-EM Data Collection, Refinement and Validation Statistics**

| | *CLN4-20 / cCpE / COP-2 / Nb*<br>*(PDB: XXXX , EMDB: XXXXX)* |
|---|---|
| **Data collection & Processing** | |
| Magnification | 120,000 |
| Voltage (kV) | 200 |
| Electron exposure (e-/Å2) | 49.4 |
| Defocus range (μm) | 0.4 - 2.0 |
| Pixel size (Å) | 0.884 |
| Symmetry imposed | C1 |
| Number of micrographs | 1302 |
| Initial particle images (no.) | 1,848,208 |
| Final particle images (no.) | 21,296 |
| Map resolution (Å)<br>FSC threshold | 4.16<br>0.143 |
| **Real-Space Refinement** | |
| Model resolution (Å)*<br>FSC threshold | 4.10 / 4.20<br>0.143 |
| Sharpening B-factor (Å2) | -74.2 |
| Dimensions (Å) | 68.95, 91.94, 152.05 |
| Chains | 5 |
| Atoms | 6652 |
| Residues | 862 |
| **B-factors (**$Å^2$) | |
| Protein | 321.89 |
| Ligand | - |

| | |
|---|---|
| Water | - |
| **R.m.s deviations** | |
| Bond lengths (Å) | 0.003 |
| Bond angles (°) | 0.781 |
| **Validation** | |
| MolProbity score | 2.35 |
| Clashscore | 21.04 |
| Poor Rotamers (%) | 0.00 |
| **Ramachandran plot** | |
| Favored (%) | 90.73 |
| Allowed (%) | 9.15 |
| Disallowed (%) | 0.12 |

* d FSC Model 0.143 Masked / Unmasked

### Supplementary Table S3| In silico success rates of the designs.

| Fold | total designs | designs passing filter | in silico success (%) |
|---|---|---|---|
| IGF | 150 | 34 | 23% |
| BBF | 72 | 26 | 36% |
| TBF | 144 | 84 | 58% |
| CLF | 750 | 52 | 7% |
| RPF | 1769 | 32 | 2% |
| GLF | 1063 | 176 | 17% |
